# Genetic Diversity Modulates The Physical And Transcriptomic Response Of Skeletal Muscle To Simulated Microgravity

**DOI:** 10.1101/2023.06.27.546810

**Authors:** Yasmina Zeineddine, Michael A. Friedman, Evan G. Buettmann, Lovell B. Abraham, Gabriel A. Hoppock, Henry J. Donahue

## Abstract

Developments in long-term space exploration necessitate advancements in countermeasures against microgravity-induced skeletal muscle loss. Astronaut data shows considerable variation in muscle loss in response to microgravity. Previous experiments suggest that genetic background influences the skeletal muscle response to unloading, but no in-depth analysis of genetic expression was performed. Here, we placed eight inbred founder strains of the diversity outbred mice (129S1/SvImJ, A/J, C57BL/6J, CAST/EiJ, NOD/ShiLtJ, NZO/HILtJ, PWK/PhJ, and WSB/EiJ) in simulated microgravity (SM) via hindlimb unloading for three weeks. Body weight, muscle morphology, muscle strength, protein synthesis marker expression, and RNA expression were collected. A/J and CAST/EiJ mice were most susceptible to SM-induced muscle loss, whereas NOD/ShiLtJ mice were the most protected. In response to SM, A/J and CAST/EiJ mice experienced reductions in body weight, muscle mass, muscle volume, and muscle cross-sectional area. A/J mice had the highest number of differentially expressed genes (68) and associated gene ontologies (328). Downregulation of immunological gene ontologies and genes encoding anabolic immune factors suggest that immune dysregulation contributes to the response of A/J mice to SM. Several muscle properties showed significant interactions between SM and mouse strain and a high degree of heritability. These data imply that genetic background plays a role in the degree of muscle loss in SM and that more individualized programs should be developed for astronauts to protect their skeletal muscles against microgravity on long term missions.

## Introduction

As the field of space exploration progresses towards missions to Mars, the development of permanent lunar bases, and the privatization of space travel, there is a concomitant need to accelerate our understanding of how to protect astronauts against the dangers of long-term missions. A primary risk of spaceflight is skeletal muscle deconditioning, which stems from the microgravity-induced unloading of the musculoskeletal system.^1, 2^ Apart from the increased risk of injury associated with losing skeletal muscle mass and strength,^3^ reloading weakened skeletal muscle fibers after exposure to microgravity can lead to injury.^4–7^ The need to transition between environments of varying gravitational field strengths without experiencing impaired locomotor function or injury emphasizes the importance of conserving skeletal muscle during exposure to microgravity.^8^ In simulated microgravity (SM), skeletal muscle mass appears to reach steady state at 70% of the preflight size after 270 days of spaceflight,^1^ and the fastest trip to Mars with current technology will take at least 7-9 months.^9^ Current nutrition and exercise interventions have been shown to attenuate microgravity-induced skeletal muscle atrophy; however, long-term missions still lead to significant reductions in muscle skeletal mass and strength,^10–15^ prompting the need for additional countermeasure strategies against muscle loss.

What is interesting about the skeletal muscle response to unloading is the range seen in the degree of muscle atrophy between individuals. Significant variability is observed between astronauts in terms of their skeletal muscle responses to spaceflight and the inter-subject response to bed rest.^12, 16–19^ Recorded volumetric losses from baseline in the skeletal muscles of the legs range between 6-24% in spaceflight and bed rest studies.^20–24^ Similarly, losses in knee extensor and plantar flexor strength have ranged from 0-29% for missions between 11 and 84 days^20, 25, 26^ and 0-55% for missions between 30-380 days, respectively.^11, 27, 28^ Within this variability, one consistent result is that the soleus, gastrocnemius, and quadriceps, known as anti-gravity muscles, are the most susceptible to unloading-induced atrophy. Their role in bipedal motion and maintenance of upright posture means that their atrophy leads to impairments in locomotion and balance upon return to stronger gravitational fields.^4, 12, 21, 29^ Determining the underlying causes of variability can enhance the development of both generic and personalized countermeasures to unloading-induced skeletal muscle atrophy, especially that of the anti-gravity muscles.

Several factors partially explain the variability in skeletal muscle response to microgravity, including preflight fitness levels and adherence to nutritional and exercise guidance during missions.^30–33^ However, the genetic component has been largely ignored. Muscle mass and strength have been shown to have high levels of heritability (phenotypic variance explained by genotype) in several studies with estimates of heritability of muscle mass ranging from 50-80% and muscle strength from 30-85%.^34^ Considering the significant genetic influence on baseline muscle mass and strength, it is reasonable to assume that genetics also severely affects how muscles respond to unloading. Judex et al. found a quantitative trait loci on chromosome 5 that was responsible for 5% of the variability in muscle cross-sectional area loss in response to unloading in mice.^35^ Maroni et al. investigated the role of genetic variability in the skeletal muscle response to disuse in founder strains of diversity outbred (DO) mice. Founder DO mice consist of eight inbred strains of mice, chosen to maximize genetic diversity and that have an extensively sequenced genome. When crossed, they form the DO mouse population, which make for ideal models for genetics-based studies.^36^ Maroni et al. placed five founder DO strains in single-hindlimb immobilization for three weeks. A mouse strain-dependent effect on disuse-induced changes in muscle mass and protein synthesis was found, suggesting that genetics play a role in the skeletal muscle response to disuse.^37^ However, no studies to date have identified specific differentially expressed genes across individuals of different genetic backgrounds that could be responsible for regulating individual responses to microgravity.

In this study, we placed the eight founder strains of DO mice into hindlimb unloading (HLU), a well-studied model of microgravity and investigated the role of genetics in both the transcriptomic and physical response of skeletal muscles to SM induced by HLU. Previous studies in our lab using these DO founder mice demonstrated that genetic variability affects the response of bone to SM.^38^ We collected data on lower limb skeletal muscle morphology, twitch force, protein synthesis markers, and RNA expression. Our hypothesis was that the morphological, biological, and transcriptomic responses of the eight founder DO strains to SM would be mouse strain-dependent. To our knowledge, this is the first study in which transcriptomic response to SM is studied using all eight strains of founder DO mice.

## Materials and Methods

### Animals

All animal procedures were completed with approval from the Virginia Commonwealth University Institutional Animal Care and Use Committee (Protocol # AD10001341). We used the eight founder strains of DO mice from the Jackson Laboratory (JAX, Bar Harbor, ME, USA: 129S1/SvImJ (stock #002448), A/J (stock #000646), C57BL/6J (stock #000664), CAST/EiJ (stock #000928), NOD/ShiLtJ (stock #001976), NZO/HILtJ (stock #002105), PWK/PhJ (stock #003715), and WSB/EiJ (stock #01145). Sixteen male mice between four and fourteen weeks old from each of the eight strains were purchased from JAX). They were given one week to acclimate to the animal facility and another week to acclimate to the wire-bottom cage environment. At sixteen weeks, half the mice from each strain were placed into SM; the other eight were left as age-matched ground controls. Mice were pair-housed in standard rat cages with wire floors. Mice were fed Teklad LM-485 chow (Envigo) and water ad libitum and kept on a 12-hour light and dark cycle. The mice in HLU were provided five pellets of chow at a time to prevent them from using their food as leverage to walk on while suspended. The duration of the study was three weeks, after which the mice were sacrificed and skeletal muscle tissue was immediately isolated for analysis.

### Simulated microgravity protocol

HLU was used as the SM model for this experiment as it unloads the hindlimbs and mimics the cephalic fluid shift that occurs in true microgravity.^39^ In addition, unlike hindlimb immobilization, HLU allows for uninhibited leg muscle contractions to occur during unloading, but no reaction forces are present to act against the muscles. The SM protocol was based off of a modified version of the Morey-Holton model.^40^ Two cross bars were added to the top of each half of the rat cages. Under general anesthesia (1-3% v/v isoflurane), the tails were wiped down with ethanol, dried, and then fully wrapped with two pieces of surgical tape. The loose ends of the tape were attached to a swivel hook attached to the end of a string that was wrapped around the center of the cross bar. The string was wound up until the animals’ hindlimbs reached an elevation of 30°. This model prevents strain on the tails of the mice while maintaining a normal load on the forelimbs, reducing stress.^41^ Since the mice were restrained due to HLU, they were unable to make contact with each other. Lab and veterinary staff inspected the condition of the mice daily. If strings were unwound from the bars, changing the angle of elevation, the mice were anaesthetized to reattach the strings properly and resuspend the mice. The body weights (BW) of the mice were measured every 7-10 days to monitor their health, but only the data from days 1 and 23 of SM were used for analysis. Mice were removed from the study if they lost more than 20% of their body weight or experienced other health issues after consultation with the veterinary staff.

### Muscle morphology

*In vivo* microCT scanning was performed to analyze changes in skeletal muscle morphology. CT has been shown to be a reliable method to assess appendicular skeletal muscle morphology.^42^ On days 1 and 23 of the procedure, mice were anaesthetized with isoflurane (2.5%) and underwent *in vivo* microCT scanning (SkyScanner 1276, Bruker, Kontich, Belgium). The animals’ hindlimbs were extended until they were straight, and then they were taped down. Their snouts were placed in a nose-cone to maintain general anesthesia by isoflurane. The scans were taken with a resolution of 60μm, exposure time of 73ms, a current of 200µA, source voltage of 60kV, and a 0.5mm aluminum filter. Average scan time of the hindlimbs lasted 10-15 seconds and resulted in less than 50mGy of radiation exposure. The scans were reconstructed using NRecon as well as the GPUReconServer reconstruction engine. Reconstructions were processed and oriented using anatomical landmarks with Dataviewer. The region of interest spanned the entire length of the tibia. Skeletal muscle cross-sectional area and volume were analyzed using CTAn with manually segmented regions of interest. Bone and subcutaneous fat were excluded from the analysis based on attenuation. Total muscle cross sectional area was analyzed and reported as the average total area of all muscles throughout the lower leg in mm^2^; total muscle volume was analyzed and reported as mm^3^.

### Muscle force testing

On day 23, the mice were placed under anesthesia with isoflurane (2.5%) and clippers were used to shave their hindlimbs. Muscle performance was measured *in vivo* using a muscle lever system (1300A system, Aurora Scientific Inc., Aurora, CAN). The mice were laid onto a heated platform and had their snouts placed in a nose-cone to maintain anesthesia. The left leg was locked into place with a knee clamp and the foot was fixed onto a 10N pedal force transducer at a 90° angle. Plantar flexion was induced through percutaneous electrical stimulation of the tibial nerve using two monopolar electrodes. Starting at 10mA, the current was increased until reaching the force response plateaued at maximum strength to find the optimal isometric twitch torque. Twitch force (under 40Hz 300ms pulse) was measured three times with five second rest periods in between. Maximum twitch force was calculated using the Aurora 605A: Dynamic Muscle Data Acquisition Analysis System. The three twitch force measurements were averaged into one value for analysis. Due to technical issues encountered during initial measurements of muscle strength, only data for C57BL/6J and A/J mice were used for analysis.

### RNA Sequencing

The gastrocnemius was extracted from the animals and incubated in RNA Later in a 4°C fridge overnight. The following day the RNA Later was removed, and the samples were stored in a −80°C freezer. A 20-30mg slice of gastrocnemius muscle was used for RNA extraction. Muscle sample total RNA was extracted with the RNeasy Fibrous Tissue Kit (Qiagen, Valencia, CA, USA) using the manufacturers’ instructions. The RNA samples were treated with DNase (Iscript gDNA Clear cDNA Synthesis Kit, Bio-Rad, Hercules, CA, USA) and tested for RNA Integrity (2100 Bioanalyzer, Agilent, Santa Clara, CA). Only samples with an RNA Integrity Number of at least 5.0 were used. DEGs with a false discovery rate (FDR) below 0.05 were used for analysis. Six samples from each strain, three control and three SM, were sequenced at the VCU Genomics Core facility using a NextSeq2000 Sequencer (Illumina). The samples were treated with the Illumina Stranded mRNA Prep to purify and fragment mRNA. Then, cDNA was synthesized, and the 3’ ends were adenylated. The samples were then treated with a ligation kit before being amplified and sequenced.

### Western Blot Analysis

Protein isolation was completed using 20mg of tissue from the quadriceps. The samples were homogenized in the buffer our lab used previously.^37^ Quantification of protein concentration was done using a BCA kit (Thermo Fisher scientific). 30μg of total protein from each sample was placed into Mini-PROTEAN (R symbol) TGX (™) 4-20% gels (Bio-Rad) and subjected to SDS-PAGE. The gels were then electroblotted onto PVDF membranes (Bio-Rad). Western blot analysis was performed to assess total and phosphorylated p70S6K1 (70kDa ribosomal protein S6 kinase) (T389, Cell Signaling Technology; Boston, MA) as well as 4EBP1 (4E-binding protein 1) (T37/46, Cell Signaling Technology). Primary antibodies were diluted at a 1:1000 ratio, and secondary antibodies were diluted 1:3000. Clarity Max (™) Western ECL Substrate (Bio-Rad) was used to detect the immune complexes. Protein quantification was completed using Image Lab software (version 6.1). To account for variation in protein loading, phosphorylated protein was normalized to total protein and the resulting ratio was used for statistical analysis.

### Statistical analysis

All statistical analyses were completed on Prism (version 9.0; GraphPad Software, La Jolla, CA). Data were analyzed using two-way ANOVAs with significance set at p<0.05 to evaluate the main effects of mouse strain, SM, and interactions. Narrow-sense heritability was calculated using one-way ANOVA, as previously described, to evaluate the additive effect of genetics on muscle properties.^37, 43^ Tukey’s multiple comparisons post-hoc test was used for data with repeated measures. Biojupies was used to perform statistical analyses on the data from RNA sequencing, including analyses for differential genes, gene ontologies, and pathway enrichment.^44^ The RNA sequencing analyses included comparing SM vs. control for each individual mouse strain in addition to combining all eight strains together and analyzing SM vs. control.

## Results

### Genetic variation affected the magnitude of changes in body weight and skeletal muscle mass

BW was analyzed on days 1 and 23 of SM (Fig. 1). There was a significant (p=0.0051) interaction between mouse strain and SM changes in BW. This interaction indicates that SM caused weight loss in some mouse strains while other mouse strains were protected from this effect. Five of the eight strains (PWK/PhJ, NZO/HILtJ, 129S1/SvImJ, CAST/EiJ, A/J) had significantly greater weight loss in the SM groups than in their control groups.

**Figure 1.**
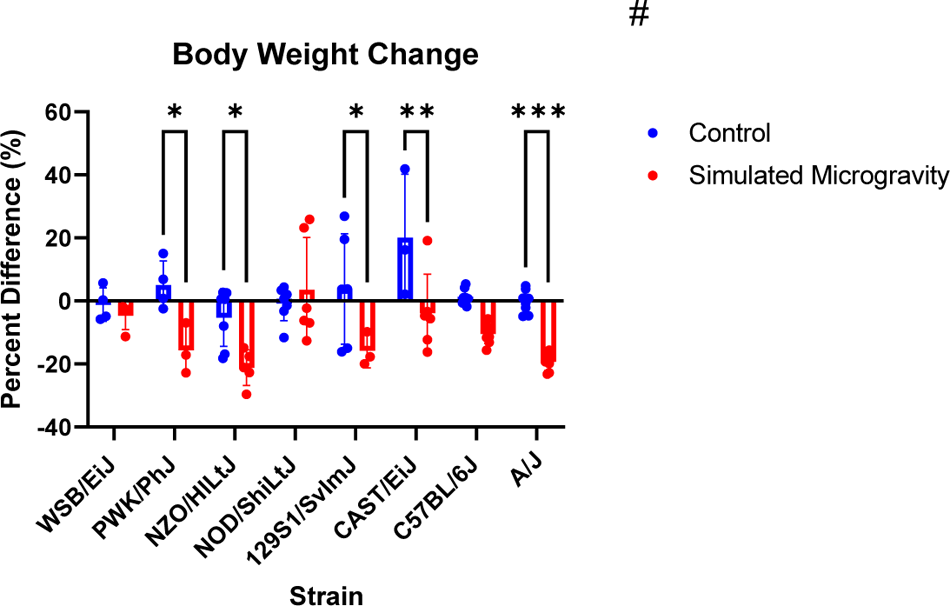
Body weight change. Change in body weight (mean ± SD) from day 1 to day 23 presented as percent difference from day 1. A/J mice demonstrated the most significant difference between the SM and control groups (p<0.001). NZO mice in SM experienced the greatest magnitude of weight loss. Analysis was performed with two-way ANOVA with Tukey’s Multiple Comparisons test (n=3-8). # represents a significant interaction between strain and unloading with p<0.05. (* p<0.05; ** p<0.01; *** p<0.001; *** p<0.0001; simulated microgravity vs control within the same strain)

Skeletal muscle masses of quadriceps and gastrocnemius were analyzed in terms of absolute mass and mass normalized by BW (Fig. 2). Baseline variation in muscle masses between strains necessitates normalizing muscle mass to BW. A significant interaction between mouse strain and SM was found for all absolute and normalized masses (quadriceps/BW: p=0.0232; gastrocnemius: p=0.015; gastrocnemius/BW: p=0.0035; soleus: p=0.0005; soleus/BW: p=0.0412), except for absolute quadriceps, which had a significant strain-dependent and unloading-dependent effect but not interaction. A/J mice demonstrated significantly lower absolute quadriceps in SM than control (p=0.0064), but this effect disappears when normalized to BW (p=0.29) as seen in Fig. 2a and Fig. 2b. WSB/EiJ mice demonstrated significantly lower quadriceps mass/BW in SM than control (p=0.0297). Both A/J and CAST/EiJ SM mice had significantly lower absolute gastrocnemius masses than their respective controls (p=0.0046; p=0.0488). These effects also disappear when masses are normalized to BW. There were no significant differences between SM and control for gastrocnemius mass/BW, absolute soleus mass, and soleus mass/BW for all mouse strains.

**Figure 2.**
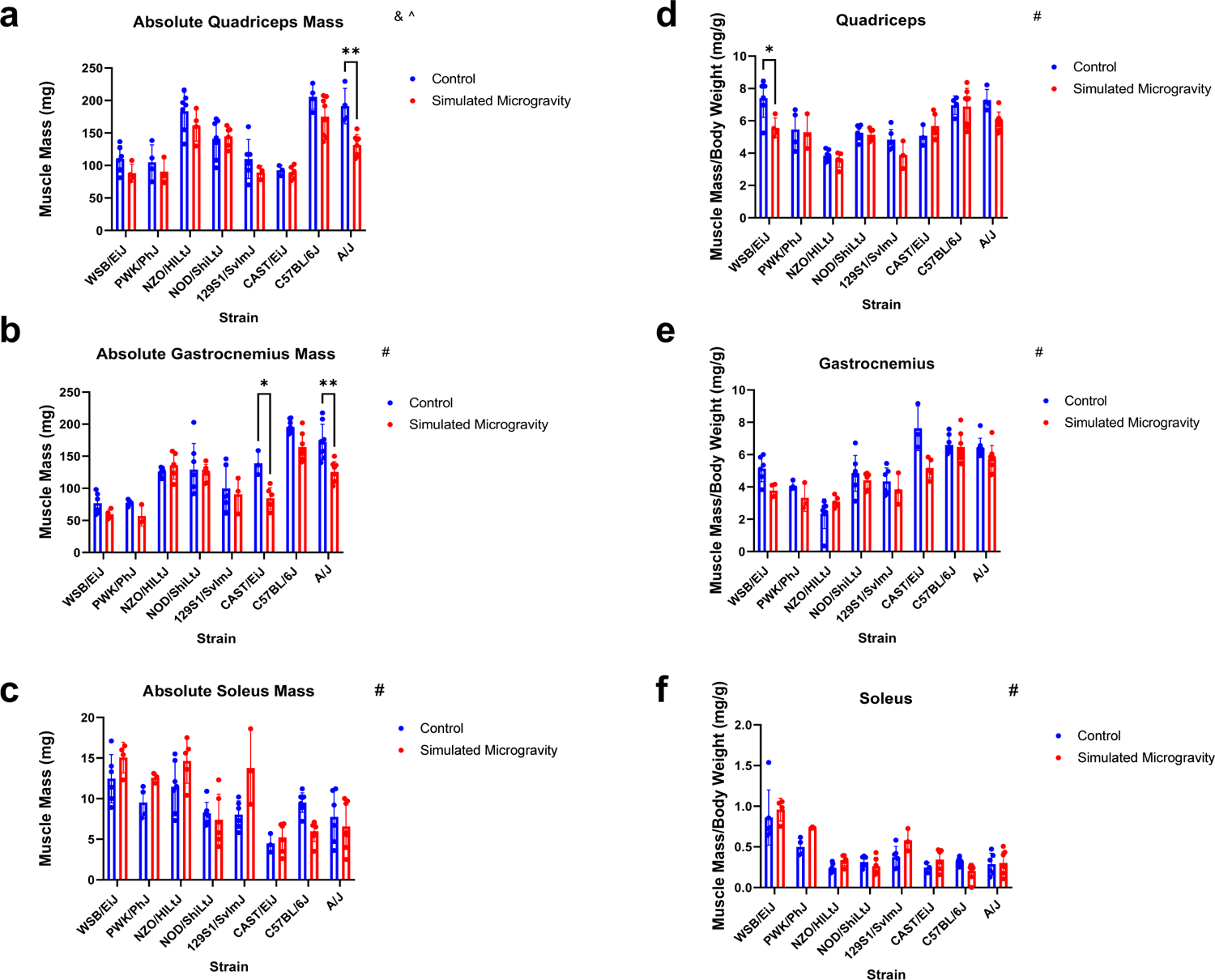
Muscle mass. (a-c): absolute wet mass in mg (mean ± SD) of the quadriceps, gastrocnemius, and soleus after three weeks in control or simulated microgravity. (d-f): muscle mass of the quadriceps, gastrocnemius, and soleus normalized to body weight (mg/g) after three weeks of control or simulated microgravity conditions. A/J mice had significantly smaller quadriceps and gastrocnemius absolute masses after exposure to simulated microgravity. CAST/EiJ mice had smaller gastrocnemius absolute masses after exposure to simulated microgravity. Analysis was performed with two-way ANOVA (n=3-8). & represents a main effect of strain; ^ represents a main effect of unloading # represents a significant interaction between strain and unloading (p<0.05). (* p<0.05; ** p<0.01; simulated microgravity vs control within the same strain)

### Cross-sectional muscle area and volume in response to SM varied by mouse strain

Cross-sectional muscle area and volume were analyzed using microCT scans of the lower limbs of the mice (Figure 3). A significant interaction between mouse strain and SM was found for both skeletal muscle cross-sectional area and volume (p=0.0009; p=0.0042). PWK/PhJ, 129S1/SvImJ, CAST/EiJ, C57BL/6J, and A/J mice exposed to SM all experienced significant losses in lower limb cross-sectional muscle area after three weeks. These same strains as well as NZO/HILtJ all experienced significant reductions in lower leg muscle volume after three weeks of SM.

**Figure 3.**
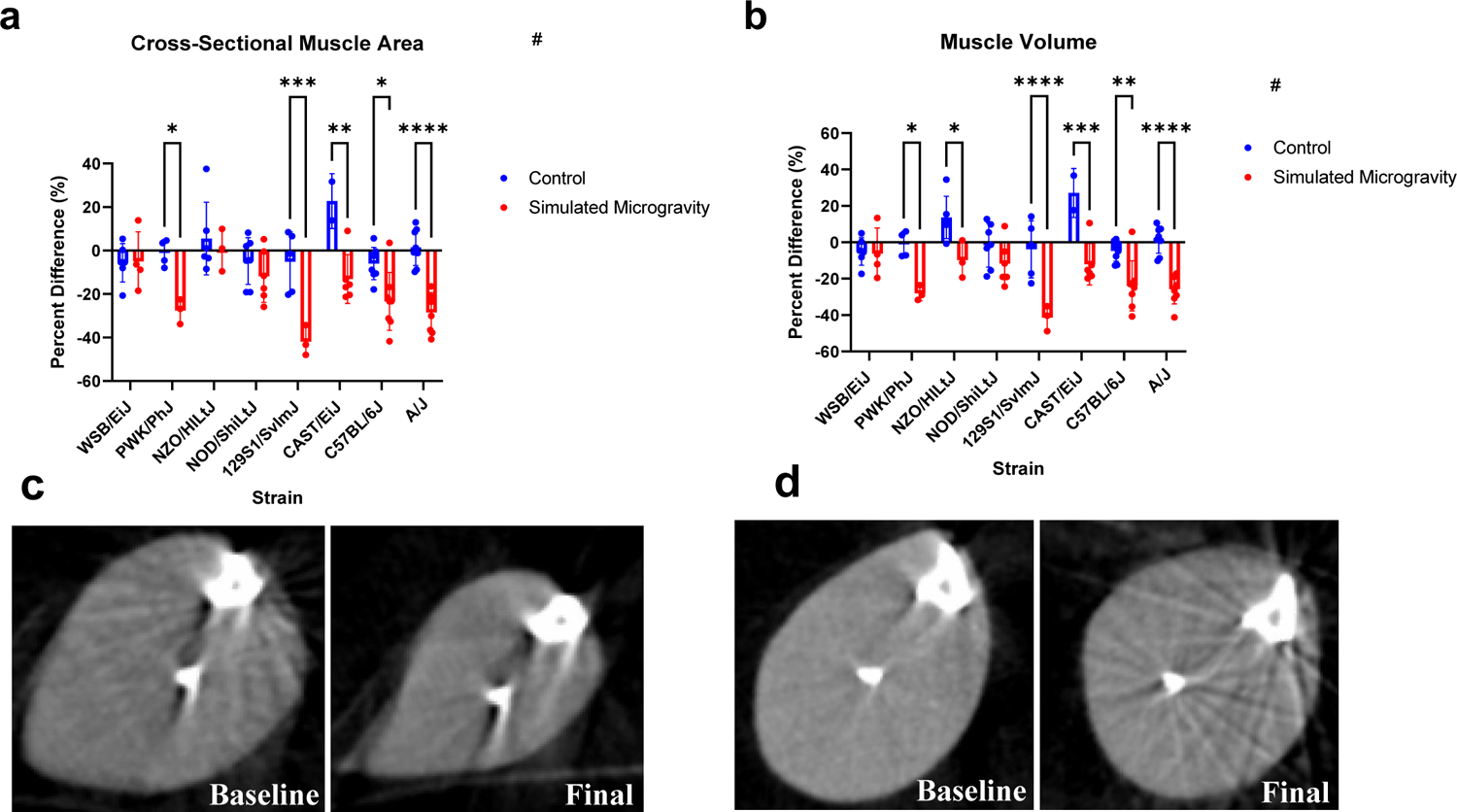
Changes in muscle cross sectional area and volume. (a): changes in lower leg cross-sectional skeletal muscle area represented as percent difference between day 1 and day 23. (b): changes in lower leg skeletal muscle volume represented as percent difference from day 1 to day 23. (c): baseline (left) and final day (right) mid-diaphyseal cross-sectional area slice from an A/J mouse. (d): baseline (left) and final day (right) mid-diaphyseal cross-sectional area slice from an NOD/ShiLtJ mouse., depicted as mean ± SD. The lower leg is defined as the segment between where the tibia and femur interface to the distal end of the tibia. A/J mice had the most significant differences in muscle volume and cross-sectional area in SM compared to control. 129S1/SvImJ mice in SM experienced the largest magnitude of loss in cross-sectional area and volume in SM compared to control. Analysis was performed with two-way ANOVA with Tukey’s post hoc test (n=3-8). # represents a significant interaction between strain and unloading (p<0.05). (* p<0.05; ** p<0.01; *** p<0.001; **** p<0.0001; simulated microgravity vs control within the same strain)

### Muscle strength

Muscle strength was analyzed using plantar flexion force measurements. Twitch force was normalized to skeletal muscle mass of the gastrocnemius to account for variation in muscle mass between strains. A significant interaction between mouse strain and SM was found for normalized twitch force (p<0.0001). For absolute twitch force, only mouse strain was found to be a significant source of variation (p<0.0001). C57BL/6J mice had significantly weaker (p<0.0001) absolute twitch forces than A/J mice in control groups. When the twitch force was normalized to skeletal muscle mass, only C57BL/6J mice displayed significantly lower forces after three weeks of SM (p<0.0001).

### RNA sequencing revealed genetic variation affected the transcriptomic response to SM

Across the eight strains there were a total of 84 differentially expressed genes (DEGs) and 935 differentially regulated gene ontologies (GOs, FDR<0.05) in response to SM (Table 1). Gene expression varied between strains following three weeks of SM. By three weeks, A/J mice had the strongest response to SM with 68 DEGs and 328 differentially regulated GOs, followed by CAST/EiJ mice, who had 11 DEGs and 99 differentially regulated GOs. NOD/ShiLtJ mice had the fewest number of DEGs (0) and differentially regulated GOs (5) in response to SM. Three genes were universally affected by SM: *Nus1, Chrnb1,* and *Dusp8* (Table 2). The most affected GO pathways in response to SM related to muscle contraction, protein transcription and translation, and metabolic processes (Table 3).

**Table 1.** Number of differentially expressed genes, gene ontologies, and enrichment pathways by strain.

| Strain | Differentially Expressed Genes |  | Differentially Expressed Gene Ontologies |  | Differentially Expressed Enrichment Pathways |  |
| --- | --- | --- | --- | --- | --- | --- |
|  | *Upregulated* | *Downregulated* | *Upregulated* | *Downregulated* | *Upregulated* | *Downregulated* |
| All strains | 1 | 2 | 8 | 51 | 0 | 54 |
| WSB/EiJ | 0 | 0 | 0 | 32 | 0 | 17 |
| PWK/PhJ | 1 | 0 | 59 | 20 | 84 | 12 |
| NZO/HILtJ | 0 | 0 | 51 | 77 | 43 | 60 |
| NOD/ShiLtJ | 0 | 0 | 0 | 5 | 0 | 1 |
| 129S1/SvImJ | 0 | 0 | 0 | 163 | 0 | 49 |
| CAST/EiJ | 8 | 3 | 69 | 30 | 16 | 32 |
| C57BL/6J | 1 | 0 | 32 | 10 | 16 | 32 |
| A/J | 57 | 11 | 30 | 298 | 36 | 163 |

**Table 2.**
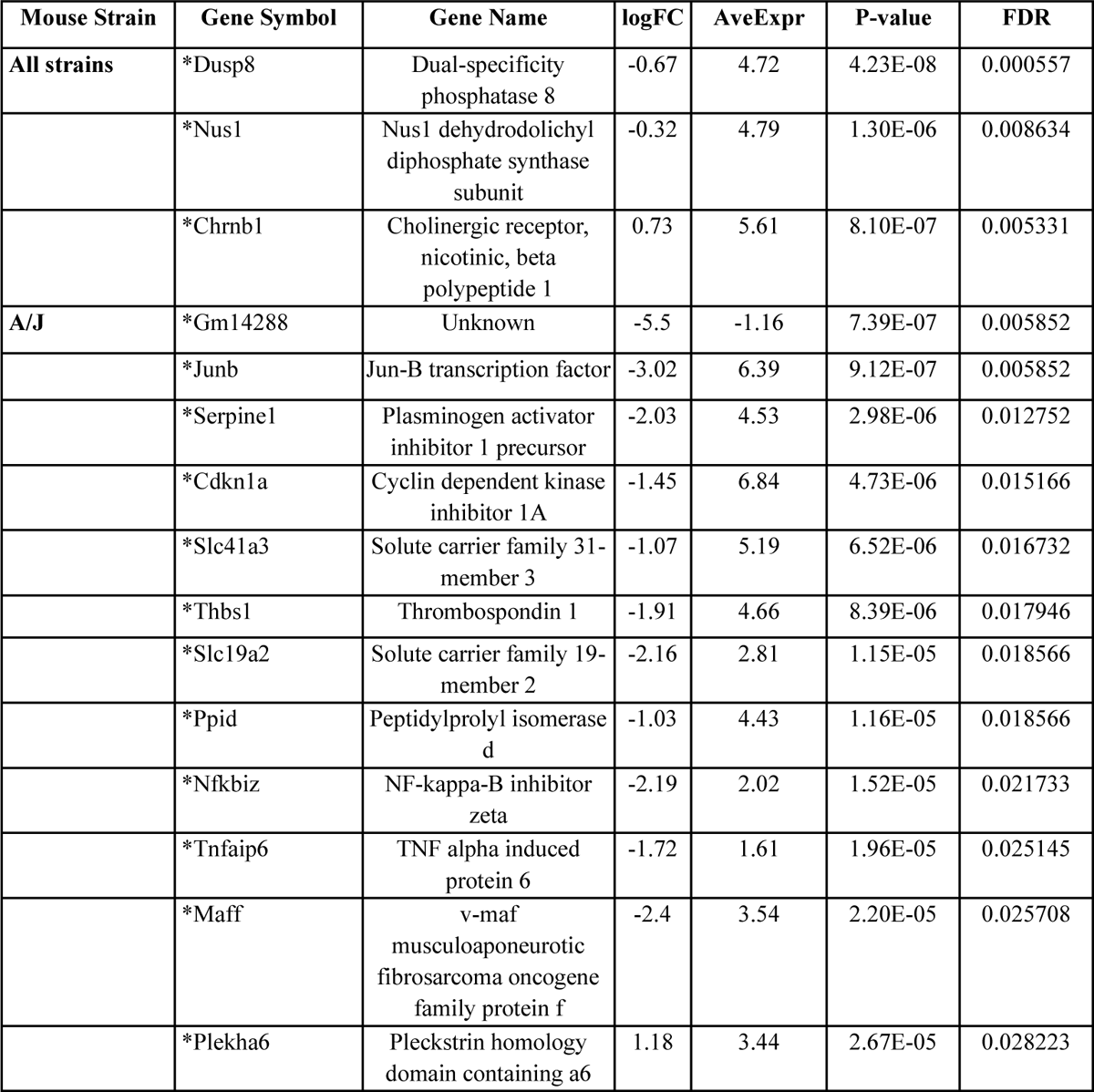
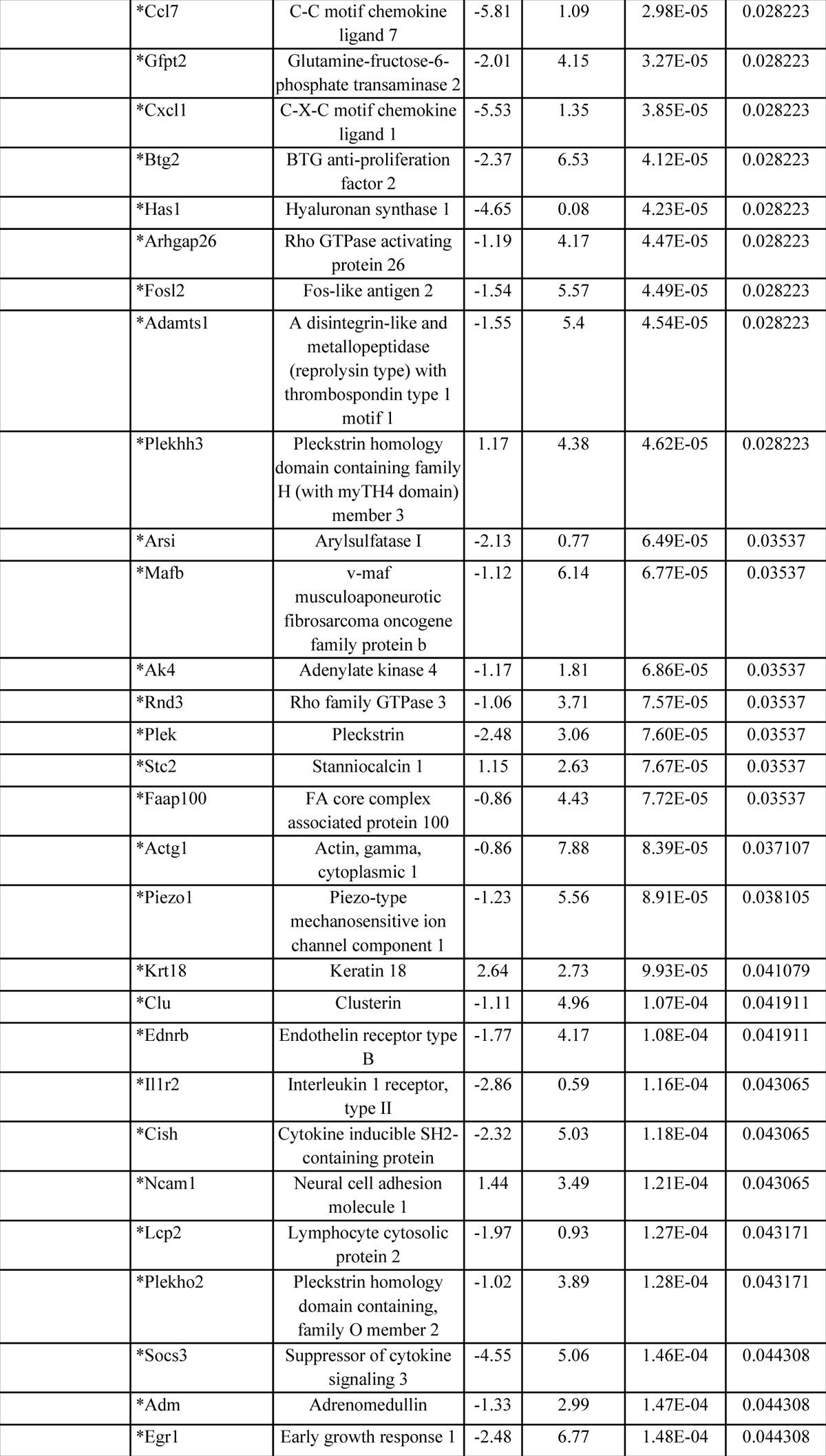
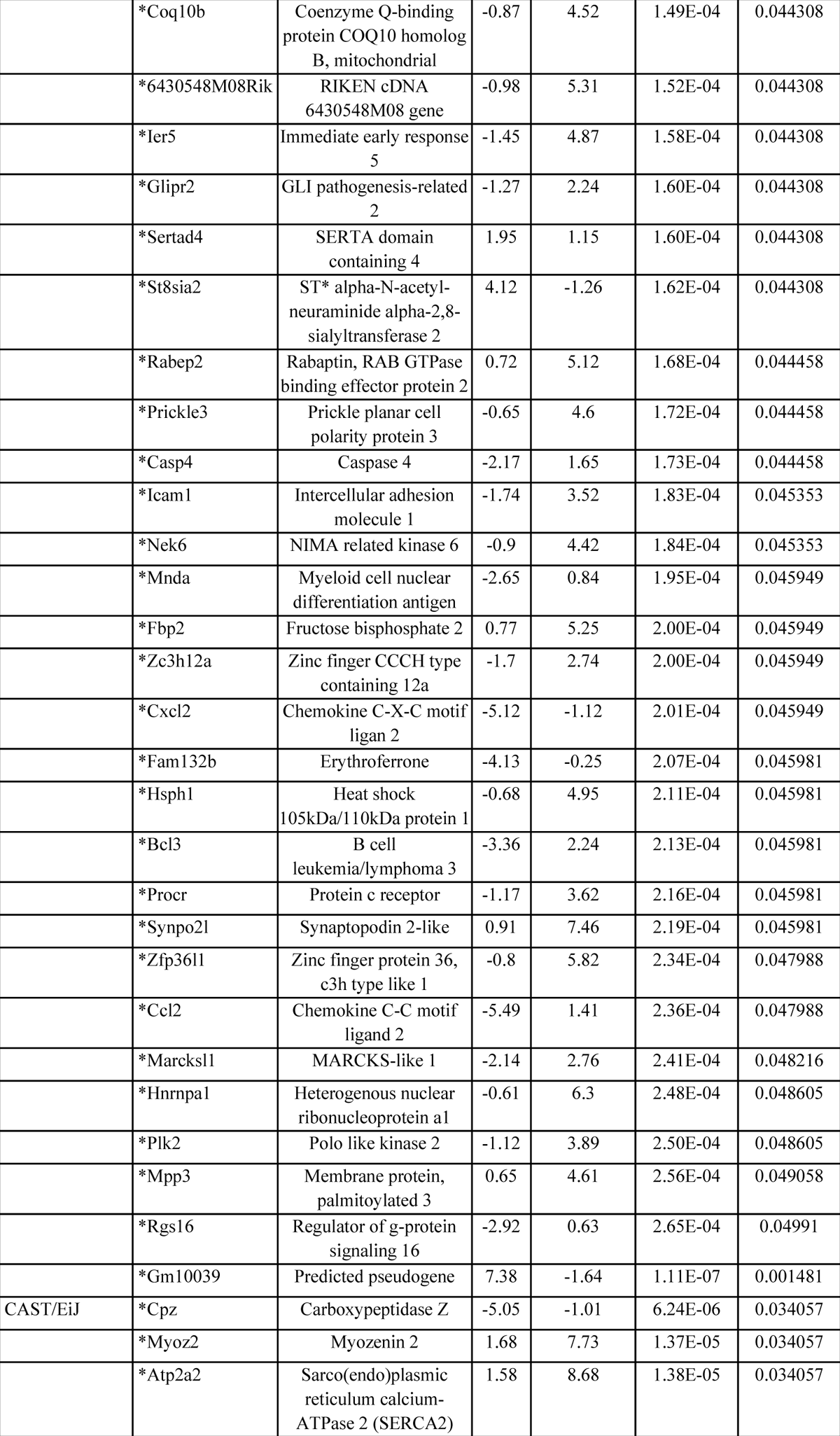
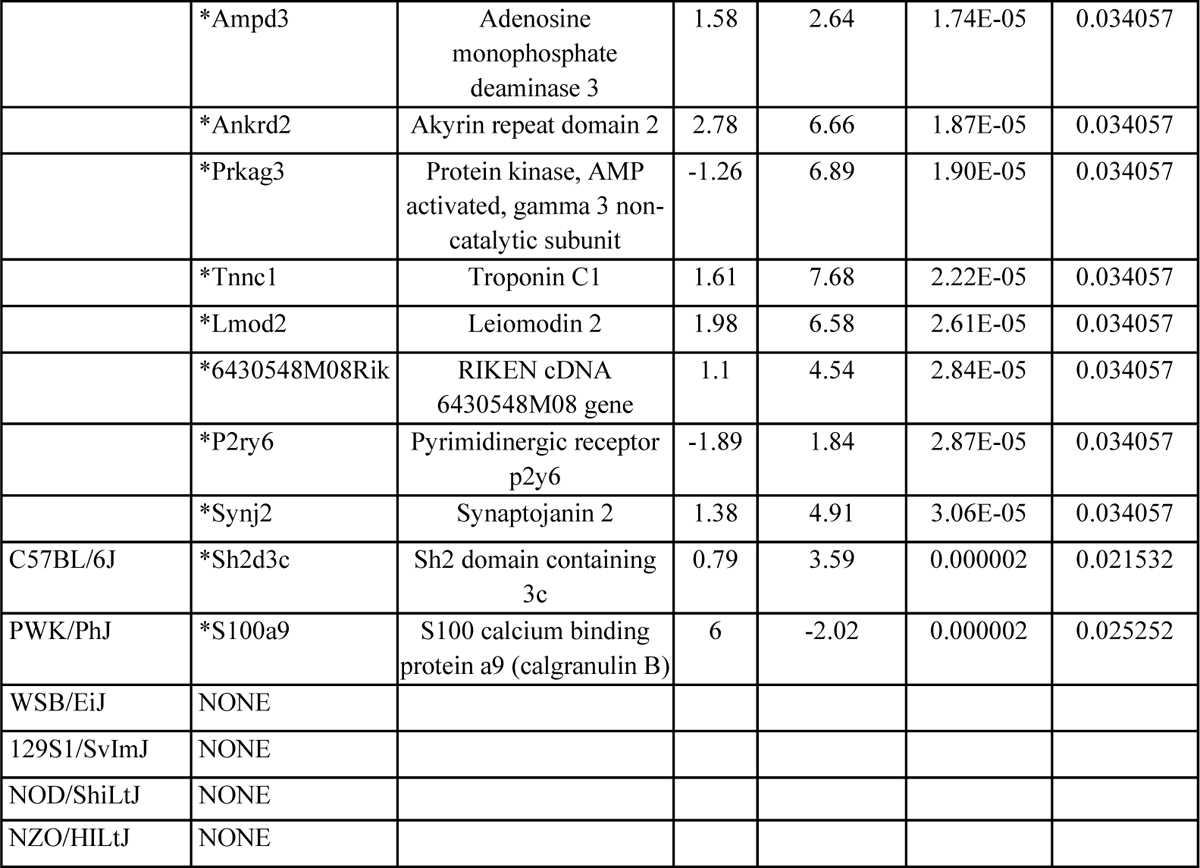
Expression and significance levels of differentially expressed genes.

**Table 3.**
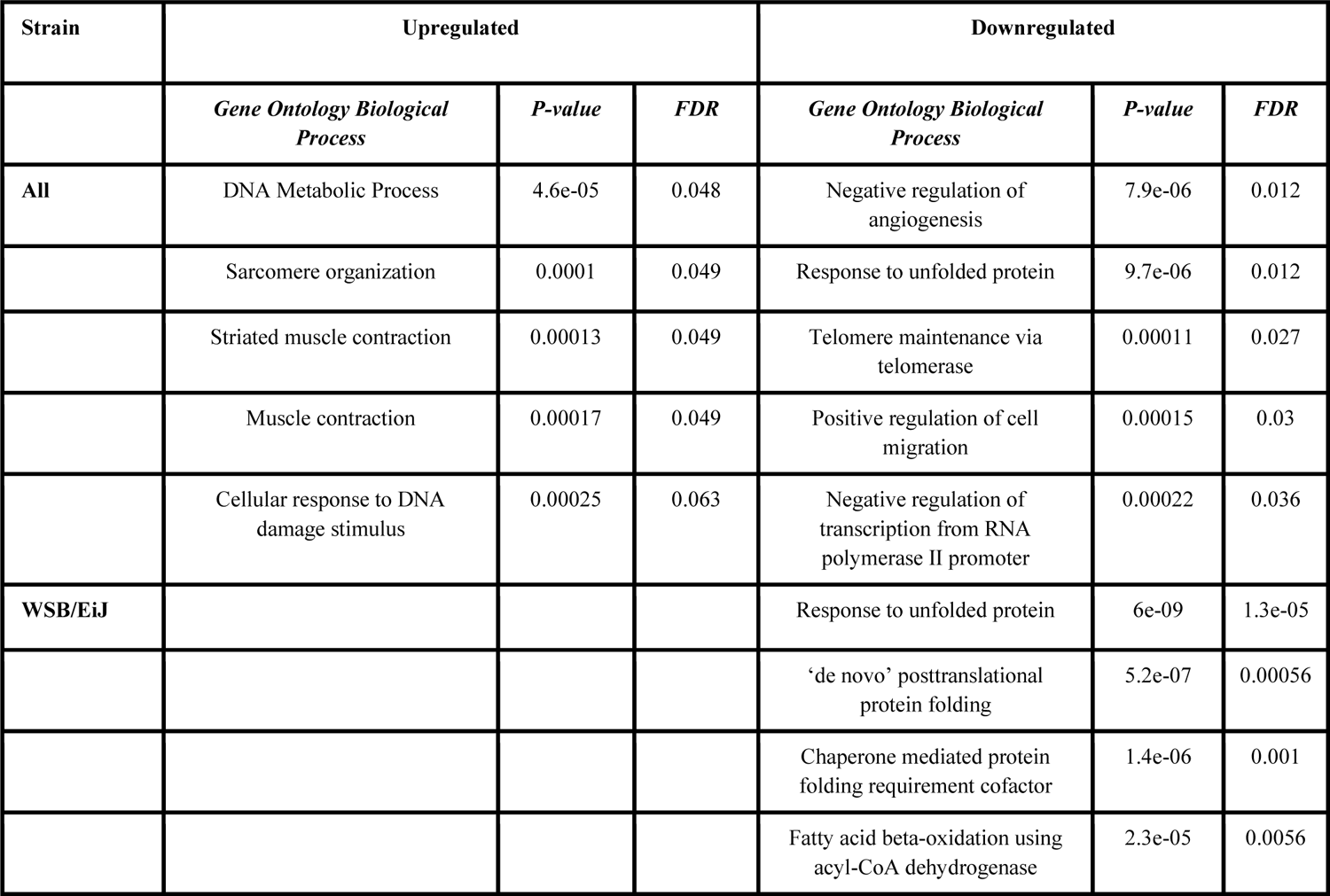
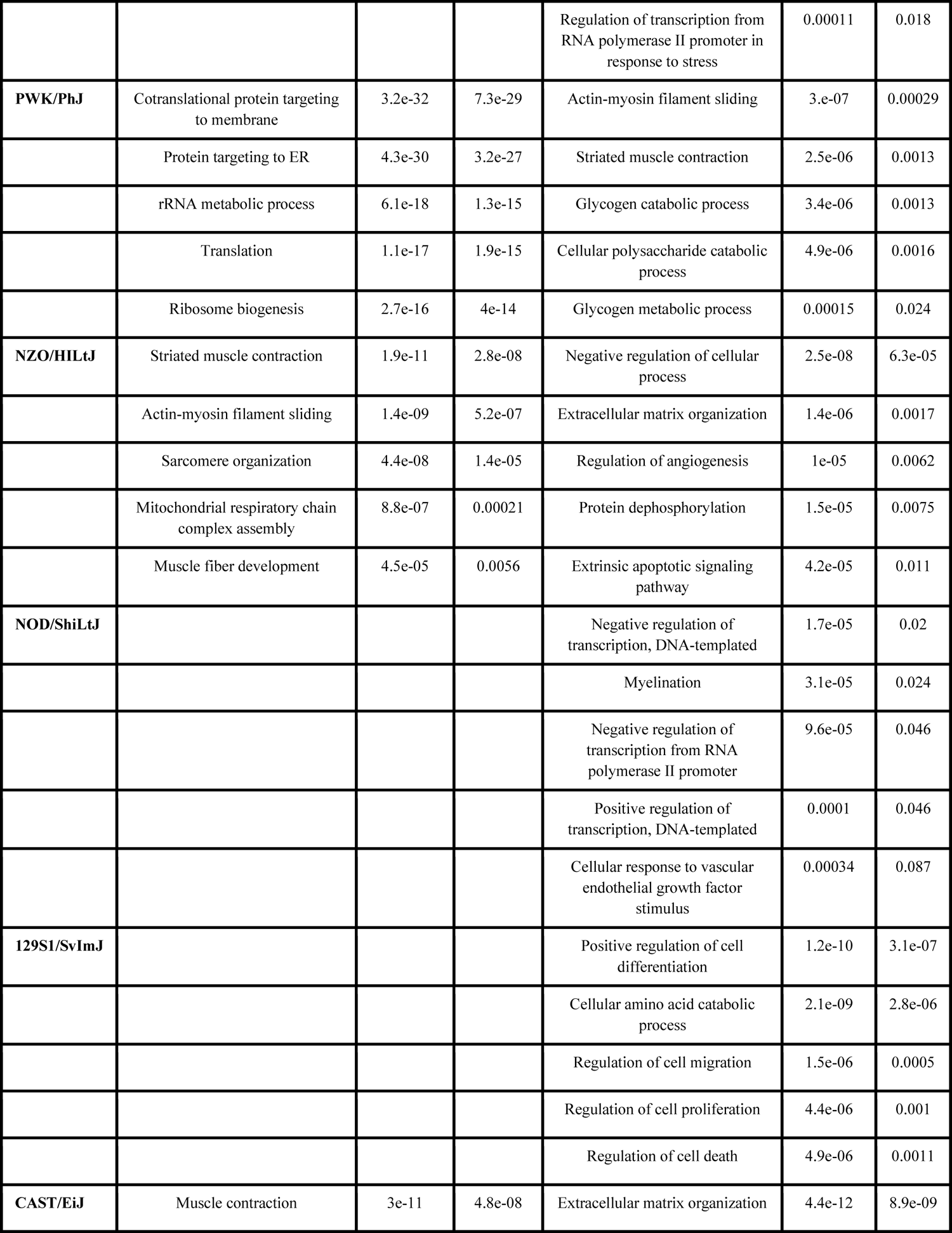
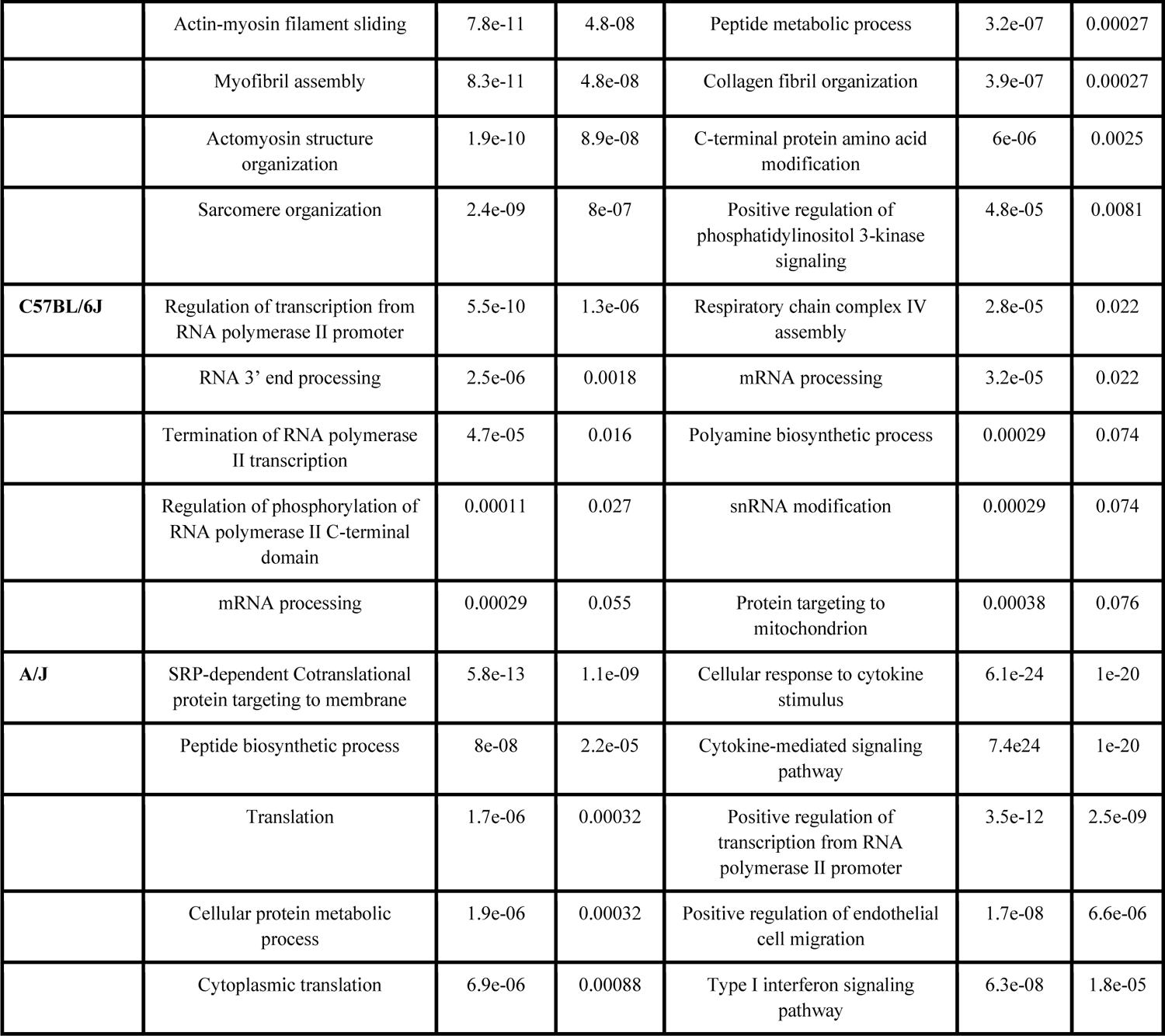
Gene ontology analysis of most upregulated pathways in simulated microgravity compared to control.

| Strain | Upregulated |  |  | Downregulated |  |  |
| --- | --- | --- | --- | --- | --- | --- |
|  | *Gene Ontology Biological Process* | *P-value* | *FDR* | *Gene Ontology Biological Process* | *P-value* | *FDR* |
| **All** | DNA Metabolic Process | 4.6e-05 | 0.048 | Negative regulation of angiogenesis | 7.9e-06 | 0.012 |
|  | Sarcomere organization | 0.0001 | 0.049 | Response to unfolded protein | 9.7e-06 | 0.012 |
|  | Striated muscle contraction | 0.00013 | 0.049 | Telomere maintenance via telomerase | 0.00011 | 0.027 |
|  | Muscle contraction | 0.00017 | 0.049 | Positive regulation of cell migration | 0.00015 | 0.03 |
|  | Cellular response to DNA damage stimulus | 0.00025 | 0.063 | Negative regulation of transcription from RNA polymerase II promoter | 0.00022 | 0.036 |
| **WSB/EiJ** |  |  |  | Response to unfolded protein | 6e-09 | 1.3e-05 |
|  |  |  |  | 'de novo' posttranslational protein folding | 5.2e-07 | 0.00056 |
|  |  |  |  | Chaperone mediated protein folding requirement cofactor | 1.4e-06 | 0.001 |
|  |  |  |  | Fatty acid beta-oxidation using acyl-CoA dehydrogenase | 2.3e-05 | 0.0056 |
|  |  |  |  | Regulation of transcription from RNA polymerase II promoter in response to stress | 0.00011 | 0.018 |
| **PWK/PhJ** | Cotranslational protein targeting to membrane | 3.2e-32 | 7.3e-29 | Actin-myosin filament sliding | 3.e-07 | 0.00029 |
|  | Protein targeting to ER | 4.3e-30 | 3.2e-27 | Striated muscle contraction | 2.5e-06 | 0.0013 |
|  | rRNA metabolic process | 6.1e-18 | 1.3e-15 | Glycogen catabolic process | 3.4e-06 | 0.0013 |
|  | Translation | 1.1e-17 | 1.9e-15 | Cellular polysaccharide catabolic process | 4.9e-06 | 0.0016 |
|  | Ribosome biogenesis | 2.7e-16 | 4e-14 | Glycogen metabolic process | 0.00015 | 0.024 |
| **NZO/HILtJ** | Striated muscle contraction | 1.9e-11 | 2.8e-08 | Negative regulation of cellular process | 2.5e-08 | 6.3e-05 |
|  | Actin-myosin filament sliding | 1.4e-09 | 5.2e-07 | Extracellular matrix organization | 1.4e-06 | 0.0017 |
|  | Sarcomere organization | 4.4e-08 | 1.4e-05 | Regulation of angiogenesis | 1e-05 | 0.0062 |
|  | Mitochondrial respiratory chain complex assembly | 8.8e-07 | 0.00021 | Protein dephosphorylation | 1.5e-05 | 0.0075 |
|  | Muscle fiber development | 4.5e-05 | 0.0056 | Extrinsic apoptotic signaling pathway | 4.2e-05 | 0.011 |
| **NOD/ShiLtJ** |  |  |  | Negative regulation of transcription, DNA-templated | 1.7e-05 | 0.02 |
|  |  |  |  | Myelination | 3.1e-05 | 0.024 |
|  |  |  |  | Negative regulation of transcription from RNA polymerase II promoter | 9.6e-05 | 0.046 |
|  |  |  |  | Positive regulation of transcription, DNA-templated | 0.0001 | 0.046 |
|  |  |  |  | Cellular response to vascular endothelial growth factor stimulus | 0.00034 | 0.087 |
| **129S1/SvImJ** |  |  |  | Positive regulation of cell differentiation | 1.2e-10 | 3.1e-07 |
|  |  |  |  | Cellular amino acid catabolic process | 2.1e-09 | 2.8e-06 |
|  |  |  |  | Regulation of cell migration | 1.5e-06 | 0.0005 |
|  |  |  |  | Regulation of cell proliferation | 4.4e-06 | 0.001 |
|  |  |  |  | Regulation of cell death | 4.9e-06 | 0.0011 |
| **CAST/EiJ** | Muscle contraction | 3e-11 | 4.8e-08 | Extracellular matrix organization | 4.4e-12 | 8.9e-09 |
|  | Actin-myosin filament sliding | 7.8e-11 | 4.8-08 | Peptide metabolic process | 3.2e-07 | 0.00027 |
|  | Myofibril assembly | 8.3e-11 | 4.8e-08 | Collagen fibril organization | 3.9e-07 | 0.00027 |
|  | Actomyosin structure organization | 1.9e-10 | 8.9e-08 | C-terminal protein amino acid modification | 6e-06 | 0.0025 |
|  | Sarcomere organization | 2.4e-09 | 8e-07 | Positive regulation of phosphatidylinositol 3-kinase signaling | 4.8e-05 | 0.0081 |
| **C57BL/6J** | Regulation of transcription from RNA polymerase II promoter | 5.5e-10 | 1.3e-06 | Respiratory chain complex IV assembly | 2.8e-05 | 0.022 |
|  | RNA 3' end processing | 2.5e-06 | 0.0018 | mRNA processing | 3.2e-05 | 0.022 |
|  | Termination of RNA polymerase II transcription | 4.7e-05 | 0.016 | Polyamine biosynthetic process | 0.00029 | 0.074 |
|  | Regulation of phosphorylation of RNA polymerase II C-terminal domain | 0.00011 | 0.027 | snRNA modification | 0.00029 | 0.074 |
|  | mRNA processing | 0.00029 | 0.055 | Protein targeting to mitochondrion | 0.00038 | 0.076 |
| **A/J** | SRP-dependent Cotranslational protein targeting to membrane | 5.8e-13 | 1.1e-09 | Cellular response to cytokine stimulus | 6.1e-24 | 1e-20 |
|  | Peptide biosynthetic process | 8e-08 | 2.2e-05 | Cytokine-mediated signaling pathway | 7.4e24 | 1e-20 |
|  | Translation | 1.7e-06 | 0.00032 | Positive regulation of transcription from RNA polymerase II promoter | 3.5e-12 | 2.5e-09 |
|  | Cellular protein metabolic process | 1.9e-06 | 0.00032 | Positive regulation of endothelial cell migration | 1.7e-08 | 6.6e-06 |
|  | Cytoplasmic translation | 6.9e-06 | 0.00088 | Type I interferon signaling pathway | 6.3e-08 | 1.8e-05 |

### p-p70S6K1 increased in response to SM in WSB mice

p70S6K1 and 4EBP1 are markers of protein synthesis within the mTOR pathway. The phosphorylation of these proteins was measured in the left quadriceps. There was a significant interaction between mouse strain and SM in p-p70S6K1/total p70S6K1 expression (p=0.002). Only WSB/EiJ mice had significantly higher (p<0.0001) p-p70S6K1/total p70S6K1 after three weeks in the SM group compared to the control group. No other strains showed differences in p-p70S6K1/total p70S6K1 between control and SM. No significant differences were found between SM and control mice in terms of p-4EBP1/total 4EBP1 expression.

### Heritability

Narrow-sense heritability, a measure of the additive effects of genetics on phenotypic variance, was calculated for several properties (Table 4). The percentages reflect the variance in the response to SM due to genetic factors. Several properties showed high levels of heritability in response to SM, including soleus/BW at 83.2%, gastrocnemius mass at 82.5%, and quadriceps mass at 78.7%. Changes in muscle area and muscle volume after SM had heritability values of 55.5% and 48.3%, respectively. In control conditions, body weight has the highest heritability at 93.8%, followed by gastrocnemius and quadriceps mass normalized to body weight. Most properties have a higher level of heritability in response to SM compared to control. Soleus muscle mass and soleus/BW heritability are much higher in response to SM (74.2%, 83.2%) than control (49.3%, 67.3%). Phosphorylation of 4EBP1 is less heritable by 22.3% in response to SM compared to control. Together, the data shows a high degree of heritability in the skeletal muscle response to SM.

**Table 4.** Heritability analysis of skeletal muscle properties.

| Property | Control | SM |
| --- | --- | --- |
| BW | 93.8% | 96.2% |
| Change in BW | 31.2% | 52.7% |
| Muscle Mass - Quadriceps | 73.5% | 78.7% |
| Muscle Mass - Gastrocnemius | 77.3% | 82.5% |
| Muscle Mass - Soleus | 49.3% | 74.2% |
| Muscle Mass/BW- Quadriceps | 77.6% | 68.6% |
| Muscle Mass/BW- Gastrocnemius | 80.2% | 77.1% |
| Muscle Mass/BW- Soleus | 67.3% | 83.2% |
| p-4EBP1 | 42.2% | 19.9% |
| p-p70S6K1 | 38.5% | 60.0% |
| Change in Muscle Area | 30.7% | 55.5% |
| Change in Muscle Volume | 44.3% | 48.3% |
| Baseline Muscle Area | 61.8% | 50.1% |
| Baseline Muscle Volume | 68.6% | 62.5% |

## Discussion

This is the first study to investigate how genetic variation affects the response to three weeks of SM in the eight founder strains of DO mice using muscle morphology and RNA sequencing data. The results suggest that genetics significantly influence the skeletal muscle response to SM, supported by variation in loss of BW, muscle size, gene expression, and muscle function by strain. A/J mice demonstrated the greatest sensitivity to SM as evidenced by the most significant differences in BW, muscle morphology, and gene expression compared to other strains. In comparison, NOD/ShiLtJ mice were most protected and had the least sensitive response to SM and demonstrated little to no responses to SM in terms of muscle morphology, protein expression, or gene expression.

BW and muscle morphology data show that there was a significant interaction between mouse strain and skeletal muscle response to SM (Figs. 1-3). A/J mice were the most susceptible strain to SM-induced muscle loss based on skeletal muscle mass and volume changes. There were no significant effects of SM on soleus muscle mass (Fig. 2). The soleus muscle is known to be highly susceptible to unloading-induced atrophy, so the lack of significant differences between SM and control in any strain was an unexpected result. However, the lower limb skeletal muscle response has only been extensively studied in C57BL/6J mice, meaning the other strains could have protective mechanisms that delay or attenuate the loss of soleus muscle following exposure to SM. Losses in muscle volume and cross-sectional area were seen in PWK/PhJ, NZO/HILtJ 129S1/SvImJ, CAST/EiJ, C57BL/6J, A/J mice and PWK/PhJ, 129S1/SvImJ, CAST/EiJ, C57BL/6J, A/J, respectively (Fig. 3). Discrepancies between muscle mass data and volume and area data can be attributed to differential responses between different muscles to SM, such as the tibialis anterior, whose individual changes could not be analyzed due to the resolution of the micro-CT scans. These data show that muscle mass and morphology as well as BW responses to SM are affected by genetics.

Decreases in postnatal muscle size during SM are typically due to atrophy of individual muscle fibers rather than decreases in the number of muscle fibers, a process mediated by imbalances in protein synthesis and degradation.^45, 46^ The degree of muscle atrophy in response to disuse has been shown to vary significantly in humans between individuals.^12, 26, 29^ The primary pathway responsible for the maintenance of skeletal muscle is mTORC1, in which mechanical load, IGF-1, or Akt act as anabolic factors that trigger downstream signaling to activate p70S6K1 and 4E-BP1 and lead to protein synthesis. Inhibition of mTORC1 leads to upregulation of MAFbx (Muscle Atrophy F-box)/Atrogin-1 and MuRF1 (Muscle Ring Finger-1), which promote protein degradation.^47^ Interestingly, despite A/J and CAST/EiJ mice demonstrating considerable losses in skeletal muscle, western blots revealed no reductions in the phosphorylation of protein synthesis markers p70S6K1 or 4EBP1 in the quadriceps. While in humans, unloading leads primarily to reduced protein synthesis, rodents have been shown to respond through early, transient reductions in protein synthesis (< 1 week following unloading), followed by increased protein degradation.^48–, 50^ This could explain why protein synthesis appeared unaffected after three weeks. It is more likely that by this stage, protein degradation would be the primary mechanism by which muscle loss occurs. In the pathway enrichment analysis, the mTOR pathway was indeed found to be downregulated in SM compared to control for all strains (FDR = 0.056). WSB/EiJ mice, who showed lower quadriceps/BW in SM compared to control, had upregulated p-p70S6K1/total p70S6K1 in SM compared to control (Fig. 4). Qualitative observations of WSB/EiJ mice in SM showed that they were more active than other mice and constantly tried to find leverage on their food pellets or water bottles for standing, which could explain the increased expression of p-p70S6K1.

**Figure 4.**
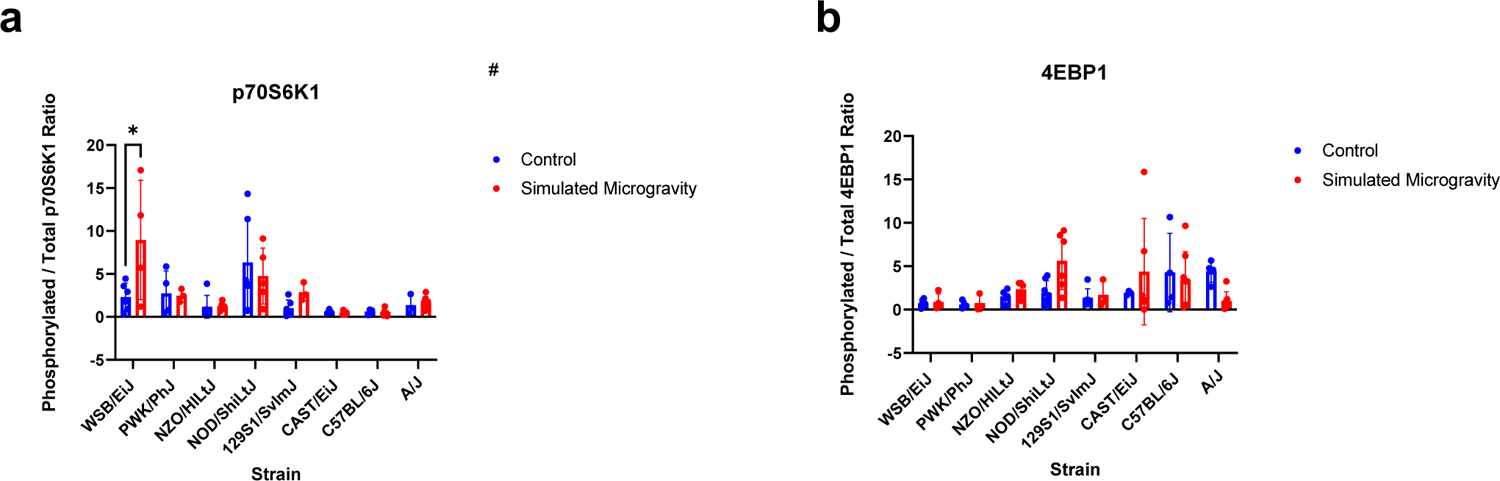
Protein synthesis markers. (A): the ratio between phosphorylated and total p70S6K1 expression (mean ± SD) in the left quadriceps of each strain after three weeks in control or simulated microgravity conditions. (B): the ratio between phosphorylated and total 4EBP1 expression (mean ± SD) in the left quadriceps of each strain after three weeks in control or simulated microgravity. Analysis was performed with two-way ANOVA (n=3-8). # represents a significant interaction between strain and unloading (p<0.05). (**** p<0.0001 simulated microgravity vs control within the same strain)

A/J mice were the most sensitive to SM, as evidenced by the greater losses in skeletal muscle mass, volume and body weight compared to other mice (Figs. 1-3), in addition to the greater number of DEGs and differentially expressed GOs (Tables 1-2). A/J mice were bred to be a model for cancer and tend to develop tumors following exposure to carcinogens. They have also been reported to have weaker responses to pathogens and are susceptible to spontaneous development of autoimmune diseases.^51, 52^ These patterns suggest that the A/J strain has a dysregulated immune response compared to other strains. Of note within the 68 DEGs in the A/J mice was the downregulation of *Piezo1, Cxcl2, Ccl2,* and *Ccl7. Piezo1* encodes a mechanosensitive ion channel known to mediate cellular responses to loading. *Piezo1* expression decreases during disuse and plays a role in the upregulation of atrophy related genes during muscle immobilization.^53^ The downregulation of *Piezo1* seen in the most SM-sensitive strain supports the association between disuse atrophy and *Piezo1* inhibition. *Cxcl2, Ccl2,* and *Ccl7* are chemokines with anabolic effects on muscle tissue.^54–59^ Their decreased expression in SM, combined with the downregulation of cytokine-mediated signaling pathway and cellular response to cytokine stimulus GOs, suggest there was suppression or alteration of immune function occurring in A/J mice exposed to SM that contributed to muscle loss. Spaceflight is known to induce immune dysregulation^60, 61^ and the immune system also plays a role in mediating the skeletal muscle response to disuse and recovery.^62–65^ The baseline immune dysfunction in A/J mice could have been exacerbated by SM and led to the particularly sensitive response of A/J to SM. This finding suggests that immune dysfunction experienced during spaceflight could increase the magnitude of skeletal muscle loss and that enhancing the strength of the immune system could attenuate skeletal muscle loss during spaceflight, though further investigation is needed to confirm this link.

Three genes were universally differentially expressed (p<0.05) in SM compared to control groups: *Dusp8* (FC of −0.67), *Chrnb1* (FC: 0.73), and *Nus1* (FC: −0.32). These three genes could be used as targets for countermeasures against unloading-induced skeletal muscle atrophy. *Dusp8* dephosphorylates mitogen-activated protein kinases, including p38, JNK, and ERK1/2. Expression of ERK1/2 is associated with the conversion from fast to slow twitch muscle fibers,^66, 67^ and its inhibition induces upregulation of Atrogin-1 and MuRF1.^68^ MuRF1 is regulated by p38 activation in HLU.^69^ Downregulation of *Dusp8* implies reduced inhibition of ERK1/2 and p38, suggesting muscle fiber type remodeling and muscle atrophy pathways were active in all SM mouse strains. *Chrnb1* encodes the beta subunit of the skeletal muscle acetylcholine receptor. Acetylcholine subunit upregulation is seen in several unloading models,^70–76^ and is indicative of neuromuscular dysfunction. However, the beta subunit positively regulates receptor turnover and metabolic half-life when phosphorylated.^77^ It is possible that its upregulation could simultaneously be a countermeasure against neuromuscular junction instability in response to SM. *Nus1* encodes the neurite outgrowth inhibitor B receptor (NgBR). NgBR is involved in several biomolecular processes, notably cholesterol trafficking,^78^ angiogenesis,^79^ neural development,^80^ and dolichol synthesis, which is required for an important and ubiquitous post-translational protein modification known as protein-N-glycosylation.^81^ Downregulation of *Nus1* could imply alterations in the rate of protein processing, decreased angiogenesis, or buildup of intracellular cholesterol deposits, processes previously reported in skeletal muscle undergoing SM.^82–86^ Together, these three DEGs show a universal response to SM irrespective of genetic background consisting of neuromuscular instability, skeletal muscle remodeling, and alterations in protein processing pathways.

GO analyses revealed further variation in transcriptomic response to SM between mouse strains (Table 2). PWK/PhJ was the only strain that demonstrated downregulations in muscle specific pathways, such as striated muscle contraction and actin-myosin filament sliding. NZO/HiLtJ and CAST/EiJ mice, on the other hand, showed upregulations in muscle contraction, actin-myosin filament sliding, and sarcomere organization, which could be protective mechanisms meant to counteract any SM-induced muscle loss. Transcription and translation-related pathways were both upregulated and downregulated across most of the strains, indicating some level of remodeling is occurring within the skeletal muscles. PWK/PhJ, C57BL/6J, and WSB/EiJ strains had downregulated metabolic pathways, including respiratory chain complex assembly and glycogen metabolism, which could be a result of reduced energy production due to decreased muscle usage. Across all strains, telomere maintenance via telomerase was downregulated, suggesting that preservation of telomeres is affected by SM regardless of genetics. Telomeres tend to be shorter after long-duration spaceflight^87, 88^ a pattern associated with age-related pathologies in humans^89^ and hematopoietic deficiencies in mice.^90^ The GO analysis show strain-dependent transcriptional responses to SM and a universal effect of SM on telomere regulation.

C57BL/6J mice showed significant losses of strength (p<0.0001) following three weeks of SM compared to those in the control group, while A/J mice showed no significant differences (Fig. 5). C57BL/6J mice showed a decrease in strength without significant changes in muscle mass. Decreases in strength are shown to precede, and often exceed, muscle atrophy during unloading, thought to be a result of neuromuscular junction degeneration, decreased neural drive to the muscles, and altered patterns of motor unit recruitment.^27, 91–93^ RNA sequencing revealed that A/J mice in SM had upregulations in *Ncam1* (FC: 1.44) and *St8sia2* (FC: 4.12)*. Ncam1* and *St8sia2* are thought to encourage neural tissue regeneration following damage by promoting neuron outgrowth and differentiation.^94–98^ These transcriptomic responses in A/J mice suggest active neuromuscular junction remodeling in the skeletal muscles of the A/J mice and potentially predict decreases in muscle strength. C57BL/6J mice showed no significant differential regulation of neuromuscular genes or pathways, suggesting that these changes may have occurred earlier in the experiment and stabilized by week three. These data show differences in the timeline of how muscle strength responds to SM between strains. Future studies should have DEG analyses from several timepoints throughout the experiment to better illustrate strain-based differences in the response to SM over time.

**Figure 5.**
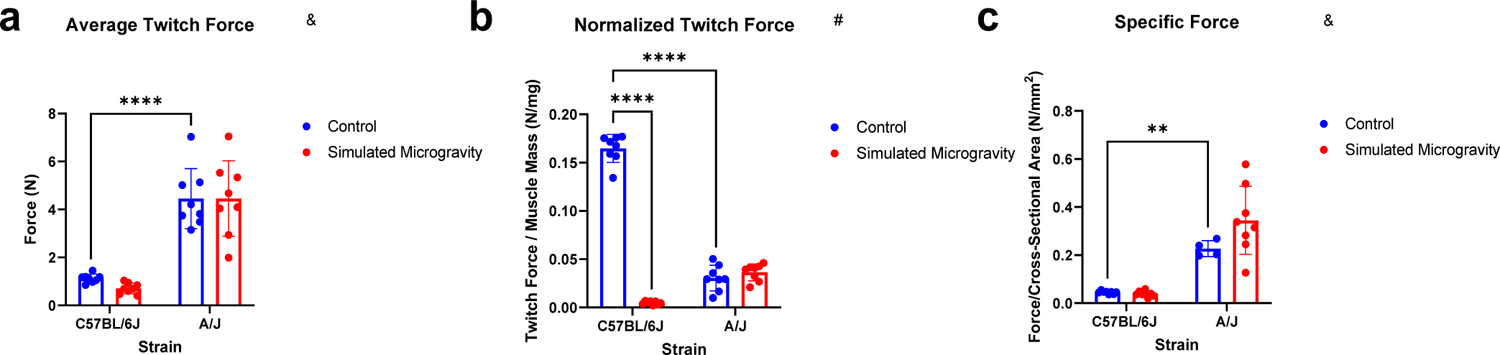
Muscle strength. (A): average absolute twitch force in C57BL/6Jand A/J mice. (B): twitch force normalized to gastrocnemius mass. (C): twitch force normalized to average cross-sectional area, also known as specific force. All forces were measured from plantar flexion induced by peripheral nerve stimulation, and data is shown as mean ± SD. 5a and 5c show baseline differences in twitch force and specific force between the C57BL/6J and A/J strains but no significant differences in response to SM within strains. 5b shows a significant loss in normalized twitch force in C57BL/6J mice in response to simulated microgravity. Analysis was performed with two-way ANOVA test (n=3-8). & represents a main effect of strain; # represents a significant interaction between strain and unloading (p<0.05). (**** p<0.0001 simulated microgravity vs control within the same strain)

Heritability analyses support and expand on previous results. For instance, we observed that change in BW and muscle mass in response to SM are highly heritable.^37^ Heritability of quadriceps/BW and gastrocnemius/BW in response to SM were 69% and 77% respectively, compared to 70% and 51% in response to single-hindlimb casting. Muscle properties in response to SM had higher levels of heritability than in control, such as soleus mass and soleus/BW. These data support the notion that mouse genetic background dictates the majority of the skeletal muscle response to SM.

A/J and CAST/EiJ strains were the most susceptible to SM while NOD/ShiLtJ and NZO/HILtJ mice were the most protected against SM-induced atrophy. NZO/HILtJ mice are an obese type II diabetic model, and NOD/ShiLtJ mice are a non-obese, type I diabetic model. These strains are prone to insulin resistance and development of diabetes as they age.^99, 100^ Type I diabetes involves a lack of insulin availability which leads to downregulation of the mTORC1 pathway.^101^ Similarly, PI3k/Akt activation is decreased in type II diabetes.^102^ These actions would be expected to accelerate disuse-atrophy in the NOD/ShiLtJ and NZO/HILtJ mice. We did not observe increased muscle atrophy in these mouse strains. However, the blood sugar levels of the mice were not measured throughout the study, and thus, we were unable to verify whether any mice fully developed diabetes. It is likely that some of the NZO/HILtJ mice were in the prediabetic or early diabetic stage as male NZO/HILtJ mice develop hyperglycemia around 8-12 weeks of age.^103^ 30-40% of NOD/HILtJ mice develop diabetes by 30 weeks,^104^ reducing the likelihood of the NOD/HILtJ mice having impaired insulin signaling during the duration of the experiment and potentially explaining the lack of expected sensitivity to SM. Further, it is possible that the HLU model we used, which increased freedom to contract the legs, protected NOD/ShiLtJ and NZO/HILtJ mice from the effects of impaired insulin signaling.

There were some limitations to this study. Our sample sizes were smaller than expected due to the stress experienced by some mouse strains in HLU (129S1/SvImJ, A/J, and NOD/HILtJ) and pair housing (PWK/PhJ and WSB/EiJ). Only male mice were used in this study to reduce the number of animals needed. However, skeletal muscle disuse atrophy is reported to be more pronounced in females compared to male mice.^105–107^ In the future, both males and females should be considered as genetic variation due to sex differences could also influence the skeletal muscle response to microgravity. Indeed, we have found this to be the case in bone response to disuse.^108^ Since the strains used in this study are inbred, they do not fully reflect the genetic diversity seen in the DO mice. Exploring how SM affects DO mice will be necessary to better understand the role of genetic variation and discover causal genes regulating the skeletal muscle response to SM.

In summary, genetics influence the skeletal muscle response to SM. For long-term missions to the Moon and Mars, countermeasure strategies to protect skeletal muscle against loss of mass and strength should take genetic background into account and be individualized for each astronaut. Further, our results suggest that more research is needed to determine the role of the immune system in mediating unloading-induced skeletal muscle atrophy, since spaceflight causes immune dysfunction and could potentially exacerbate the skeletal muscle loss seen in astronauts. Future studies should use DO mice to better understand the specific genes and alleles that regulate the skeletal muscle response to unloading and identify protective genes that could assist the development of interventions to prevent unloading-induced muscle atrophy. Potential gene targets identified in this study include *Dusp8, Chrbn1,* and *Nus1*.

## Supporting information

Gene Expression Data

Mouse Data

Western Blots

## Author Contributions

YZ – Writing: original draft, review and editing; Investigation; Visualization; Formal Analysis. MAF – Conceptualization; Writing – review and editing; Data curation; Investigation; Methodology; Formal Analysis; Supervision. EGB – Writing: review and editing; Investigation. LBA – Investigation. GAH – Investigation. HJD – Conceptualization; Funding Acquisition; Methodology; Supervision; Writing: review and editing.

## Acknowledgements

The RNA sequencing data included in this study was generated at the Genomics Core facility at Virginia Commonwealth University. This work was supported by NASA grant 80NSSC18K1473, the Alice T. and William H. Goodwin Jr. Research Endowment, and the Translational Research Institute for Space Health Postdoctoral Fellowship (NASA Cooperative Agreement NNX16AO69A). The funder played no role in study design, data collection, analysis and interpretation of data, or the writing of this manuscript.

## Data Availability

All data generated or analyzed during this study are included in this published article and its supplementary information files.

## References

1. di Prampero, P. E. & Narici, M. V. Muscles in microgravity: from fibres to human motion. J. Biomech. 36, 403–412 (2003).

2. Comfort, P. et al. Effects of Spaceflight on Musculoskeletal Health: A Systematic Review and Meta-analysis, Considerations for Interplanetary Travel. Sports Med. Auckl. Nz 51, 2097–2114 (2021).

3. Yeung, S. S. Y. et al. Sarcopenia and its association with falls and fractures in older adults: A systematic review and meta-analysis. J. Cachexia Sarcopenia Muscle 10, 485–500 (2019).

4. Fitts, R. H., Riley, D. J. & Widrick, J. J. Physiology of a Microgravity Environment Invited Review: Microgravity and skeletal muscle. https://journals.physiology.org/doi/full/10.1152/jappl.2000.89.2.823?rfr_dat=cr_pub++0pubmed&url_ver=Z39.88-2003&rfr_id=ori%3Arid%3Acrossref.org (2000).

5. Riley, D. A. et al. In-flight and postflight changes in skeletal muscles of SLS-1 and SLS-2 spaceflown rats. J. Appl. Physiol. Bethesda Md 1985 81, 133–144 (1996).

6. Riley, D. A. et al. Muscle sarcomere lesions and thrombosis after spaceflight and suspension unloading. J. Appl. Physiol. Bethesda Md 1985 73, 33S–43S (1992).

7. Riley, D. A. et al. Decreased thin filament density and length in human atrophic soleus muscle fibers after spaceflight. J. Appl. Physiol. Bethesda Md 1985 88, 567–572 (2000).

8. Lane, H. W. & Schoeller, D. A. Nutrition in Spaceflight and Weightlessness Models. (CRC Press, 1999).

9. Levchenko, I., Xu, S., Mazouffre, S., Keidar, M. & Bazaka, K. Mars Colonization: Beyond Getting There. Glob. Chall. 3, 1800062 (2018).

10. Bergouignan, A. et al. Towards human exploration of space: The THESEUS review series on nutrition and metabolism research priorities. NPJ Microgravity 2, 16029 (2016).

11. Antonutto, G., Capelli, C., Girardis, M., Zamparo, P. & di Prampero, P. E. Effects of microgravity on maximal power of lower limbs during very short efforts in humans. J. Appl. Physiol. Bethesda Md 1985 86, 85–92 (1999).

12. Trappe, S. et al. Exercise in space: human skeletal muscle after 6 months aboard the International Space Station. J. Appl. Physiol. Bethesda Md 1985 106, 1159–1168 (2009).

13. Momken, I. et al. Resveratrol prevents the wasting disorders of mechanical unloading by acting as a physical exercise mimetic in the rat. FASEB J. Off. Publ. Fed. Am. Soc. Exp. Biol. 25, 3646–3660 (2011).

14. Servais, S., Letexier, D., Favier, R., Duchamp, C. & Desplanches, D. Prevention of unloading-induced atrophy by vitamin E supplementation: links between oxidative stress and soleus muscle proteolysis? Free Radic. Biol. Med. 42, 627–635 (2007).

15. English, K. L. et al. High intensity training during spaceflight: results from the NASA Sprint Study. Npj Microgravity 6, 1–9 (2020).

16. Böcker, J., Schmitz, M.-T., Mittag, U., Jordan, J. & Rittweger, J. Between-Subject and Within-Subject Variaton of Muscle Atrophy and Bone Loss in Response to Experimental Bed Rest. Front. Physiol. 12, 743876 (2022).

17. Fernandez-Gonzalo, R. et al. Substantial and Reproducible Individual Variability in Skeletal Muscle Outcomes in the Cross-Over Designed Planica Bed Rest Program. Front. Physiol. 12, (2021).

18. Fitts, R. H. et al. Prolonged space flight-induced alterations in the structure and function of human skeletal muscle fibres. J. Physiol. 588, 3567–3592 (2010).

19. Zange, J. et al. Changes in calf muscle performance, energy metabolism, and muscle volume caused by long-term stay on space station MIR. Int. J. Sports Med. 18 **Suppl 4**, S308–309 (1997).

20. Akima, H. et al. Effect of short-duration spaceflight on thigh and leg muscle volume. Med. Sci. Sports Exerc. 32, 1743–1747 (2000).

21. Narici, M. V. & de Boer, M. D. Disuse of the musculo-skeletal system in space and on earth. Eur. J. Appl. Physiol. 111, 403–420 (2011).

22. LeBlanc, A. et al. Muscle volume, MRI relaxation times (T2), and body composition after spaceflight. J. Appl. Physiol. 89, 2158–2164 (2000).

23. Gopalakrishnan, R. et al. Muscle Volume, Strength, Endurance, and Exercise Loads During 6-Month Missions in Space. Aviat. Space Environ. Med. 81, 91–104 (2010).

24. LeBlanc, A., Rowe, R., Schneider, V., Evans, H. & Hedrick, T. Regional muscle loss after short duration spaceflight. Aviat. Space Environ. Med. 66, 1151–1154 (1995).

25. Tesch, P. A., Berg, H. E., Bring, D., Evans, H. J. & LeBlanc, A. D. Effects of 17-day spaceflight on knee extensor muscle function and size. Eur. J. Appl. Physiol. 93, 463–468 (2005).

26. Trappe, S. W. et al. Comparison of a space shuttle flight (STS-78) and bed rest on human muscle function. J. Appl. Physiol. Bethesda Md 1985 91, 57–64 (2001).

27. Lambertz, D., Pérot, C., Kaspranski, R. & Goubel, F. Effects of long-term spaceflight on mechanical properties of muscles in humans. J. Appl. Physiol. 90, 179–188 (2001).

28. Koryak, Y. A. Architectural and functional specifics of the human triceps surae muscle in vivo and its adaptation to microgravity. J. Appl. Physiol. Bethesda Md 1985 126, 880–893 (2019).

29. Fitts, R. H., Riley, D. R. & Widrick, J. J. Functional and structural adaptations of skeletal muscle to microgravity. J. Exp. Biol. 204, 3201–3208 (2001).

30. Moosavi, D., et al. The Effects of Spaceflight Microgravity on the Musculoskeletal System of Humans and Animals, with an Emphasis on Exercise as a Countermeasure: A Systematic Scoping Review. Physiol. Res. 70, 119–151 (2021).

31. Lee, P. H. U., Chung, M., Ren, Z., Mair, D. B. & Kim, D.-H. Factors mediating spaceflight-induced skeletal muscle atrophy. Am. J. Physiol.-Cell Physiol. (2022) doi:10.1152/ajpcell.00203.2021.

32. Greenisen, M. C., Hayes, J. C. & Siconolfi, S. F. Functional Performance Evaluation. (1999).

33. Scott-Conner, C. et al. Review of NASA’s Evidence Reports on Human Health Risks: 2015 Letter Report. (2016).

34. Roth, S. M. Genetic aspects of skeletal muscle strength and mass with relevance to sarcopenia. BoneKEy Rep. 1, 58 (2012).

35. Judex, S., Zhang, W., Donahue, L. R. & Ozcivici, E. Genetic and tissue level muscle-bone interactions during unloading and reambulation. J. Musculoskelet. Neuronal Interact. 16, 174–182 (2016).

36. Svenson, K. L. et al. High-Resolution Genetic Mapping Using the Mouse Diversity Outbred Population. Genetics 190, 437–447 (2012).

37. Maroni, C. R. et al. Genetic variability affects the response of skeletal muscle to disuse. J. Musculoskelet. Neuronal Interact. 21, 387–396 (2021).

38. Friedman, M. A. et al. Genetic Variation affects the Response to Hindlimb Suspension in Bones of the Founder Strains of the Diversity Outbred Mice. (Under Review). JOR (2023).

39. Juhl, O. J. et al. Update on the effects of microgravity on the musculoskeletal system. NPJ Microgravity 7, 28 (2021).

40. Morey-Holton, E. R. & Globus, R. K. Hindlimb unloading rodent model: technical aspects. J. Appl. Physiol. 92, 1367–1377 (2002).

41. Hargens, A. R., Steskal, J., Johansson, C. & Tipton, C. M. Tissue fluid shift, forelimb loading, and tail tension in tail-suspended rats. Physiol. Suppl. 27, (1984).

42. Mitsiopoulos, N. et al. Cadaver validation of skeletal muscle measurement by magnetic resonance imaging and computerized tomography. J. Appl. Physiol. 85, 115–122 (1998).

43. Lariviere, W. R. & Mogil, J. S. The Genetics of Pain and Analgesia in Laboratory Animals. Methods Mol. Biol. Clifton NJ 617, 261–278 (2010).

44. Torre, D., Lachmann, A. & Ma’yan, A. BioJupies: Automated Generation of Interactive Notebooks for RNA-Seq Data Analysis in the Cloud. Cell Systems. https://maayanlab.cloud/biojupies/ (2018).

45. Atherton, P. J. & Smith, K. Muscle protein synthesis in response to nutrition and exercise. J. Physiol. 590, 1049–1057 (2012).

46. Schiaffino, S., Dyar, K. A., Ciciliot, S., Blaauw, B. & Sandri, M. Mechanisms regulating skeletal muscle growth and atrophy. FEBS J. 280, 4294–4314 (2013).

47. Bodine, S. C. et al. Identification of ubiquitin ligases required for skeletal muscle atrophy. Science 294, 1704–1708 (2001).

48. Mirzoev, T., Tyganov, S., Vilchinskaya, N., Lomonosova, Y. & Shenkman, B. Key Markers of mTORC1-Dependent and mTORC1-Independent Signaling Pathways Regulating Protein Synthesis in Rat Soleus Muscle During Early Stages of Hindlimb Unloading. Cell. Physiol. Biochem. 39, 1011– 1020 (2016).

49. Krawiec, B. J., Frost, R. A., Vary, T. C., Jefferson, L. S. & Lang, C. H. Hindlimb casting decreases muscle mass in part by proteasome-dependent proteolysis but independent of protein synthesis. Am. J. Physiol. Endocrinol. Metab. 289, E969–980 (2005).

50. de Boer, M. D. et al. The temporal responses of protein synthesis, gene expression and cell signalling in human quadriceps muscle and patellar tendon to disuse. J. Physiol. 585, 241–251 (2007).

51. Kang, T. J., Lee, G. S., Kim, S. K., Jin, S. H. & Chae, G. T. Comparison of Two Mice Strains, A/J and C57BL/6, in Caspase-1 Activity and IL-1<svg style=“vertical-align:-4.698pt;width:13.15px;” id=“M1” height=“21.9375” version=“1.1” viewBox=“0 0 13.15 21.9375” width=“13.15” xmlns:xlink=“http://www.w3.org/1999/xlink” xmlns=“http://www.w3.org/2000/svg”> <g transform=“matrix(.022,-0,0,-.022,.062,16.025)”><path id=“x1D6FD” d=“M558 587q0-32-14-61t-40-53.5t-48.5-41t-54.5-36.5q144-51144-174q0-55-43.5-108t-104.5-87q-77-42-131-42q-310-5420t-3147l1118q48-29108-29q790119.543t40.5109t-44.5107.5t-119.550.5l2247q3416521q966196157q042-2467.5t-6225.5q-240-43.5-9t-35-29.5t-27-44t-22.5-63t-19.5-75.5t-18.5-91q-57-294-68-380q-26-190-35-200q-26-31-97-37l-426q19948170l77413q2312152.5187.5t83.5114.5q706214862q51088.5-34t37.5-91z”/></g> </svg> Secretion of Macrophage to <i >Mycobacterium leprae Infection. Mediators Inflamm. 2010, e708713 (2010).

52. Sellers, R. S., Clifford, C. B., Treuting, P. M. & Brayton, C. Immunological Variation Between Inbred Laboratory Mouse Strains: Points to Consider in Phenotyping Genetically Immunomodified Mice. Vet. Pathol. 49, 32–43 (2012).

53. Hirata, Y. et al. A Piezo1/KLF15/IL-6 axis mediates immobilization-induced muscle atrophy. J. Clin. Invest. 132, e154611.

54. Shireman, P. K. et al. MCP-1 deficiency causes altered inflammation with impaired skeletal muscle regeneration. J. Leukoc. Biol. 81, 775–785 (2007).

55. Lu, H., Huang, D., Ransohoff, R. M. & Zhou, L. Acute skeletal muscle injury: CCL2 expression by both monocytes and injured muscle is required for repair. FASEB J. 25, 3344–3355 (2011).

56. Shireman, P. K., Contreras-Shannon, V., Reyes-Reyna, S. M., Robinson, S. C. & McManus, L. M. MCP-1 parallels inflammatory and regenerative responses in ischemic muscle. J. Surg. Res. 134, 145–157 (2006).

57. Little, H. C. et al. Multiplex Quantification Identifies Novel Exercise-regulated Myokines/Cytokines in Plasma and in Glycolytic and Oxidative Skeletal Muscle*. Mol. Cell. Proteomics 17, 1546–1563 (2018).

58. Huang, P. et al. Feasibility, potency, and safety of growing human mesenchymal stem cells in space for clinical application. NPJ Microgravity 6, 16 (2020).

59. Kitase, Y. et al. CCL7 Is a Protective Factor Secreted by Mechanically Loaded Osteocytes. J. Dent. Res. 93, 1108–1115 (2014).

60. Crucian, B. & Choukér, A. Immune System in Space: General Introduction and Observations on Stress-Sensitive Regulations. in *Stress Challenges and Immunity in Space: From Mechanisms to Monitoring and Preventive Strategies* (ed. Choukèr, A.) 205–220 (Springer International Publishing, 2020). doi:10.1007/978-3-030-16996-1_11.

61. Crucian, B. E. et al. Immune System Dysregulation During Spaceflight: Potential Countermeasures for Deep Space Exploration Missions. Front. Immunol. 9, 1437 (2018).

62. Teodori, L., Costa, A., Campanella, L. & Albertini, M. C. Skeletal Muscle Atrophy in Simulated Microgravity Might Be Triggered by Immune-Related microRNAs. Front. Physiol. 9, (2019).

63. You, Z. et al. Molecular feature of neutrophils in immune microenvironment of muscle atrophy. J. Cell. Mol. Med. 26, 4658–4665 (2022).

64. Dumont, N. & Frenette, J. Macrophages Protect against Muscle Atrophy and Promote Muscle Recovery in Vivo and in Vitro. Am. J. Pathol. 176, 2228–2235 (2010).

65. Reidy, P. T., Dupont-Versteegden, E. E. & Drummond, M. J. Macrophage Regulation of Muscle Regrowth from Disuse in Aging. Exerc. Sport Sci. Rev. 47, 246–250 (2019).

66. Liu, R. et al. DUSP8 Regulates Cardiac Ventricular Remodeling by Altering ERK1/2 Signaling. Circ. Res. 119, 249–260 (2016).

67. Boyer, J. G., et al. ERK1/2 signaling induces skeletal muscle slow fiber-type switching and reduces muscular dystrophy disease severity. JCI Insight 4, (2019).

68. Shi, H. et al. Mitogen-activated protein kinase signaling is necessary for the maintenance of skeletal muscle mass. Am. J. Physiol. Cell Physiol. 296, C1040–1048 (2009).

69. Kim, J. et al. p38 MAPK Participates in Muscle-Specific RING Finger 1-Mediated Atrophy in Cast-Immobilized Rat Gastrocnemius Muscle. Korean J. Physiol. Pharmacol. Off. J. Korean Physiol. Soc. Korean Soc. Pharmacol. 13, 491–496 (2009).

70. Agrin-Induced Phosphorylation of the Acetylcholine Receptor Regulates Cytoskeletal Anchoring and Clustering - PMC. https://www.ncbi.nlm.nih.gov/pmc/articles/PMC2185523/.

71. Friese, M. B., Blagden, C. S. & Burden, S. J. Synaptic differentiation is defective in mice lacking acetylcholine receptor β-subunit tyrosine phosphorylation. Development 134, 4167–4176 (2007).

72. Yanez, P. & Martyn, J. A. J. Prolonged d-Tubocurarine Infusion and/or Immobilization Cause Upregulation of Acetylcholine Receptors and Hyperkalemia to Succinylcholine in Rats. Anesthesiology 84, 384–391. (1996).

73. Khan, M. A. S. et al. Non-surgically-induced disuse muscle atrophy and neuromuscular dysfunction upregulates alpha7 acetylcholine receptors. Can. J. Physiol. Pharmacol. 92, 1–8 (2014).

74. Gomes, R. R. & Booth, F. W. Expression of acetylcholine receptor mRNAs in atrophying and nonatrophying skeletal muscles of old rats. J. Appl. Physiol. 85, 1903–1908 (1998).

75. Wang, H., Yang, B., Han, G. & Li, S. Potency of nondepolarizing muscle relaxants on muscle-type acetylcholine receptors in denervated mouse skeletal muscle. Acta Pharmacol. Sin. 31, 1541–1546 (2010).

76. Hogue, C. W., Itani, M. S. & Martyn, J. A. Resistance to d-tubocurarine in lower motor neuron injury is related to increased acetylcholine receptors at the neuromuscular junction. Anesthesiology 73, 703–709 (1990).

77. Rudell, J. B. & Ferns, M. J. Regulation of muscle acetylcholine receptor turnover by β subunit tyrosine phosphorylation. Dev. Neurobiol. 73, 399–410 (2013).

78. Harrison, K. D. et al. Nogo-B Receptor stabilizes Niemann-Pick Type C2 protein and regulates intracellular cholesterol trafficking. Cell Metab. 10, 208–218 (2009).

79. Miao, R. Q. et al. Identification of a receptor necessary for Nogo-B stimulated chemotaxis and morphogenesis of endothelial cells. Proc. Natl. Acad. Sci. U. S. A. 103, 10997–11002 (2006).

80. Eckharter, C. et al. Schwann Cell Expressed Nogo-B Modulates Axonal Branching of Adult Sensory Neurons Through the Nogo-B Receptor NgBR. Front. Cell. Neurosci. 9, 454 (2015).

81. Harrison, K. D. et al. Nogo-B receptor is necessary for cellular dolichol biosynthesis and protein N-glycosylation. EMBO J. 30, 2490–2500 (2011).

82. Pagano, A. F. et al. Short-term disuse promotes fatty acid infiltration into skeletal muscle. J. Cachexia Sarcopenia Muscle 9, 335–347 (2018).

83. Gorgey, A. S. & Dudley, G. A. Skeletal muscle atrophy and increased intramuscular fat after incomplete spinal cord injury. Spinal Cord 45, 304–309 (2007).

84. Gordon, B. S., Kelleher, A. R. & Kimball, S. R. Regulation of muscle protein synthesis and the effects of catabolic states. Int. J. Biochem. Cell Biol. 45, 2147–2157 (2013).

85. Delp, M. D., Colleran, P. N., Wilkerson, M. K., McCurdy, M. R. & Muller-Delp, J. Structural and functional remodeling of skeletal muscle microvasculature is induced by simulated microgravity. Am. J. Physiol.-Heart Circ. Physiol. 278, H1866–H1873 (2000).

86. Colleran, P. N. et al. Alterations in skeletal perfusion with simulated microgravity: a possible mechanism for bone remodeling. J. Appl. Physiol. 89, 1046–1054 (2000).

87. Luxton, J. J. & Bailey, S. M. Twins, Telomeres, and Aging-in Space! Plast. Reconstr. Surg. 147, 7S–14S (2021).

88. Luxton, J. J. et al. Telomere Length Dynamics and DNA Damage Responses Associated with Long-Duration Spaceflight. Cell Rep. 33, 108457 (2020).

89. Vaiserman, A. & Krasnienkov, D. Telomere Length as a Marker of Biological Age: State-of-the-Art, Open Issues, and Future Perspectives. Front. Genet. 11, (2021).

90. Calado, R. T. & Young, N. S. Telomere Diseases. N. Engl. J. Med. 361, 2353–2365 (2009).

91. Alford, E. K., Roy, R. R., Hodgson, J. A. & Edgerton, V. R. Electromyography of rat soleus, medical gastrocnemius, and tibialis anterior during hind limb suspension. Exp. Neurol. 96, 635–649 (1987).

92. Edgerton, V. R. & Roy, R. R. Neuromuscular adaptation to actual and simulated weightlessness. Adv. Space Biol. Med. 4, 33–67 (1994).

93. Ohira, T., Kawano, F., Goto, K., Kaji, H. & Ohira, Y. Responses of neuromuscular properties to unloading and potential countermeasures during space exploration missions. Neurosci. Biobehav. Rev. 136, 104617 (2022).

94. Szewczyk, L. M. et al. ST8SIA2 promotes oligodendrocyte differentiation and the integrity of myelin and axons. Glia 65, 34–49 (2017).

95. Werneburg, S. et al. Polysialylation at Early Stages of Oligodendrocyte Differentiation Promotes Myelin Repair. J. Neurosci. 37, 8131–8141 (2017).

96. Monti, E. et al. Neuromuscular junction instability and altered intracellular calcium handling as early determinants of force loss during unloading in humans. J. Physiol. 599, 3037–3061 (2021).

97. Murgia, M. et al. Signatures of muscle disuse in spaceflight and bed rest revealed by single muscle fiber proteomics. PNAS Nexus 1, pgac086 (2022).

98. Chipman, P. H., Schachner, M. & Rafuse, V. F. Presynaptic NCAM Is Required for Motor Neurons to Functionally Expand Their Peripheral Field of Innervation in Partially Denervated Muscles. J. Neurosci. 34, 10497–10510 (2014).

99. The Jackson Laboratory. 001976 - NOD Strain Details. https://www.jax.org/strain/001976.

100. The Jackson Laboratory. 002105 - New Zealand Obese Strain Details. The Jackson Laboratory https://www.jax.org/strain/002105.

101. Mitch, W. E. & Goldberg, A. L. Mechanisms of Muscle Wasting — The Role of the Ubiquitin– Proteasome Pathway. N. Engl. J. Med. 335, 1897–1905 (1996).

102. Workeneh, B. & Bajaj, M. The regulation of muscle protein turnover in diabetes. Int. J. Biochem. Cell Biol. 45, 2239–2244 (2013).

103. Crofford, O. B. & Davis, C. K. GROWTH CHARACTERISTICS, GLUCOSE TOLERANCE AND INSULIN SENSITIVITY OF NEW ZEALAND OBESE MICE. Metabolism. 14, 271–280 (1965).

104. Chen, D., Thayer, T. C., Wen, L. & Wong, F. S. Mouse Models of Autoimmune Diabetes: The Nonobese Diabetic (NOD) Mouse. Methods Mol. Biol. Clifton NJ 2128, 87–92 (2020).

105. Shackelford, L. C. et al. Resistance exercise as a countermeasure to disuse-induced bone loss. J. Appl. Physiol. Bethesda Md 1985 97, 119–129 (2004).

106. Deschenes, M. R., McCoy, R. W. & Mangis, K. A. Factors Relating to Gender Specificity of Unloading-induced Declines in Strength. Muscle Nerve 46, 210–217 (2012).

107. Rosa-Caldwell, M. E. et al. Female mice may have exacerbated catabolic signalling response compared to male mice during development and progression of disuse atrophy. J. Cachexia Sarcopenia Muscle 12, 717–730 (2021).

108. DeNapoli, R. C., Buettmann, E. G., Friedman, M. A., Lichtman, A. H. & Donahue, H. J. Global cannabinoid receptor 1 deficiency affects disuse-induced bone loss in a site-specific and sex-dependent manner. J. Biomech. 146, 111414 (2023).

