## Supplementary material for "Genetic Diversity Modulates The Physical And Transcriptomic Response Of Skeletal Muscle To Simulated Microgravity": Western Blots

**Western Blot Supplementary Images** Original Western Blot Images. Western blots were performed to quantify levels of protein synthesis markers p70S6K1 (70kDa) and 4EBP1 (15kDa). SM indicates a sample that was in simulated microgravity; Con indicates a control sample. N/A indicates a sample that was not part of the groups of interest and was not included in the data. Samples displaying an "X" beneath their lane indicate that they were not used for quantification due to quality issues n=3-8.

**Supplementary Figure 1.** PWK/PhJ p70S6K1 original western blot images. Quantification of total and phosphorylated p70S6K1 protein levels in PWK/PhJ mouse quadriceps. (a): total p70S6K1. SM n=3; Con n=4. (b): phosphorylated p70S6K1 (T-389). SM n=3; Con n=4.

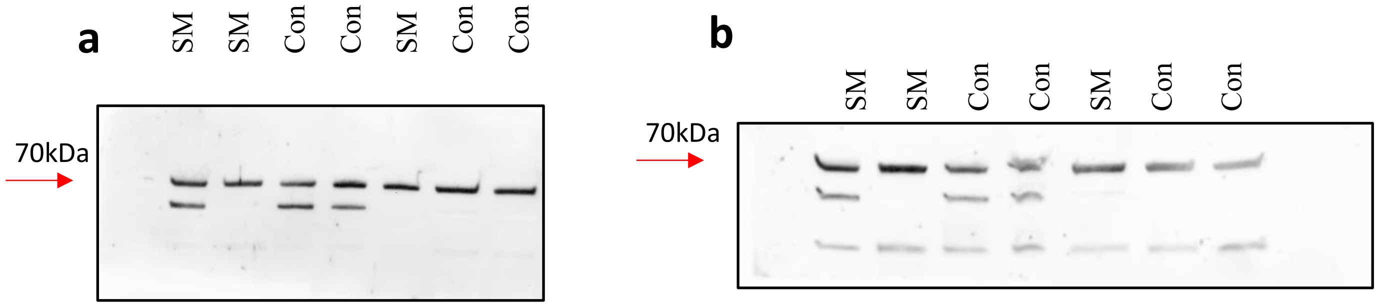


**Supplementary Figure 2.** PWK/PhJ 4EBP1 original western blot images. Quantification of total and phosphorylated 4EBP1 protein levels in PWK/PhJ mouse quadriceps. n = 7. (a): total 4EBP1. SM n=3; Con n=4. (b): phosphorylated 4EBP1 (T37/46). SM n=3; Con n=4.

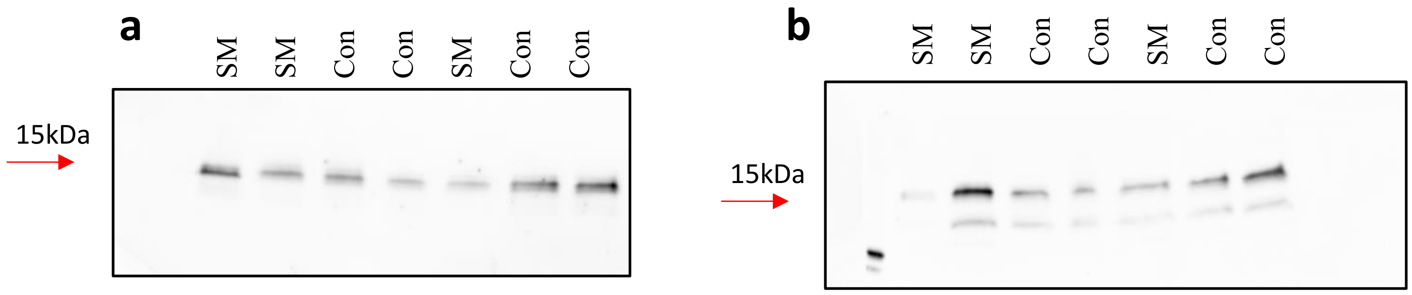

**Supplementary Figure 3.** WSB/EiJ p70S6K1 original western blot images. Quantification of total and phosphorylated p70S6K1 protein levels in WSB/EiJ mouse quadriceps. (a): total p70S6K1. The final sample was run on a gel separate from that on the left due to limitations in space. SM n=4; Con n=10. (b): phosphorylated p70S6K1 (T-389).

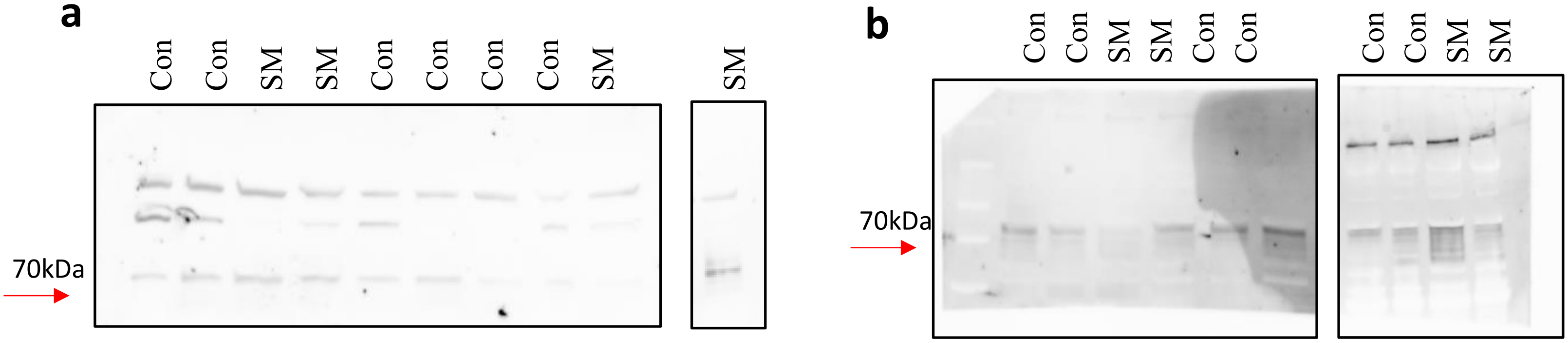


**Supplementary Figure 4.** WSB/EiJ 4EBP1 original western blot images. Quantification of total and phosphorylated 4EBP1 protein levels in WSB/EiJ mouse quadriceps. (a): total 4EBP1. The final sample was run on a gel separate from that on the left due to limitations in space. SM n=4; Con n=10. (b): phosphorylated 4EBP1 (T37/46). SM n=4; Con n=10.

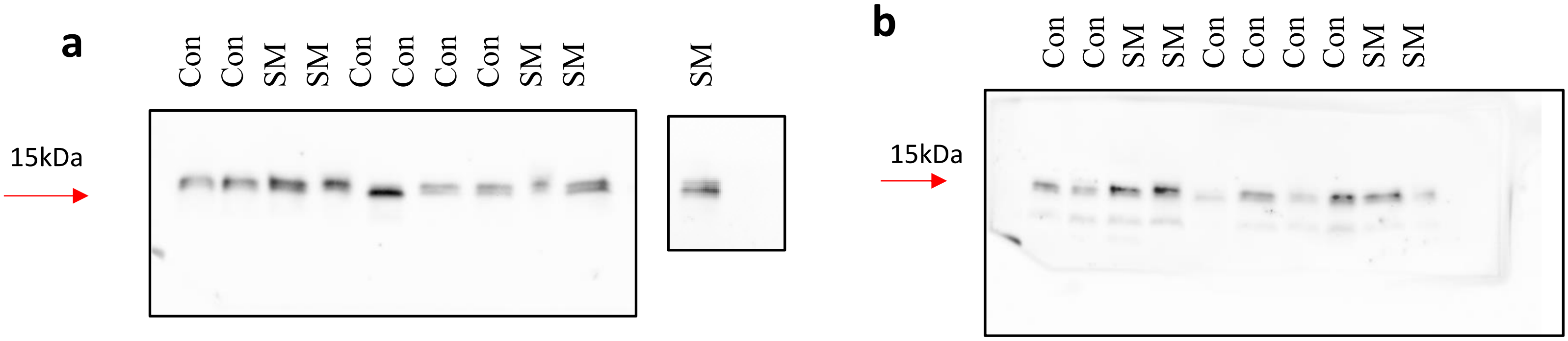

**Supplementary Figure 5.** CAST/EiJ p70S6K1 original western blot images. Quantification of total and phosphorylated p70S6K1 protein levels in CAST/EiJ mouse quadriceps. (a): total p70S6K1. SM n=3; Con n=6. (b): phosphorylated p70S6K1 (T-389). SM n=3; Con n=6.

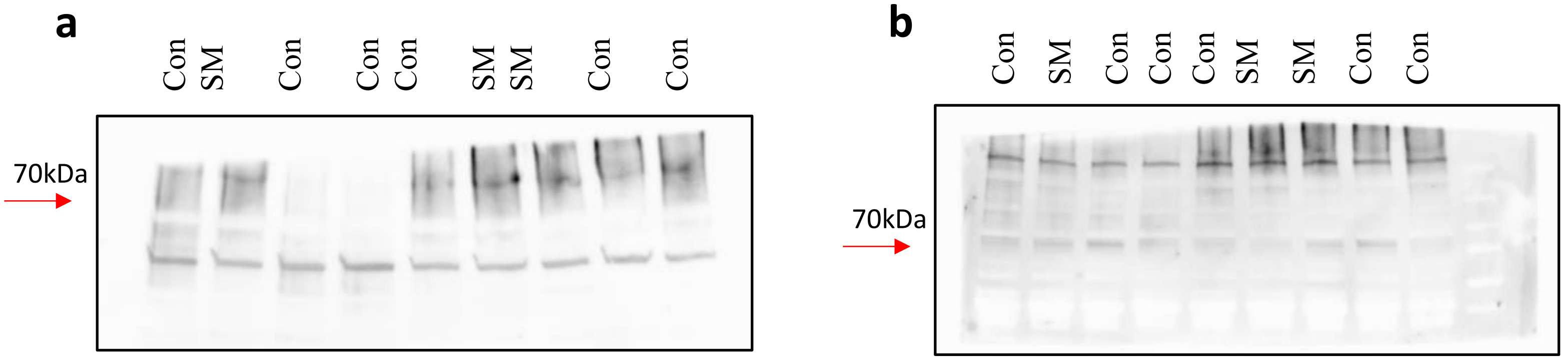


**Supplementary Figure 6.** CAST/EiJ 4EBP1 original western blot images. Quantification of total and phosphorylated 4EBP1 protein levels in CAST/EiJ mouse quadriceps. (a): total 4EBP1. SM n=3; Con n=6. (b): phosphorylated 4EBP1 (T37/46). SM n=3; Con n=6.

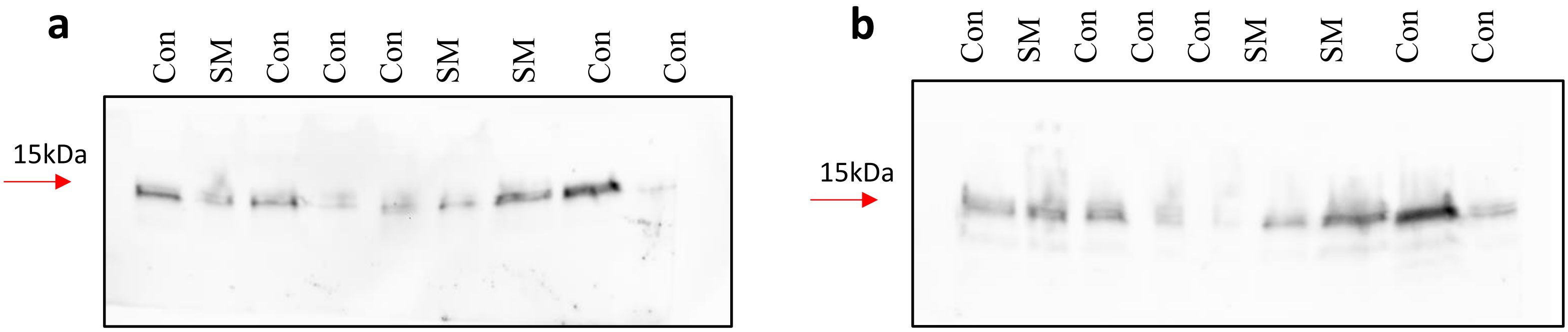

**Supplementary Figure 7.** 129S1/SvImJ p70S6K1 original western blot images. Quantification of total and phosphorylated p70S6K1 protein levels in 129S1/SvImJ mouse quadriceps. (a): total p70S6K1. SM n=3; Con n=6. (b): phosphorylated p70S6K1 (T-389). SM n=3; Con n=6.

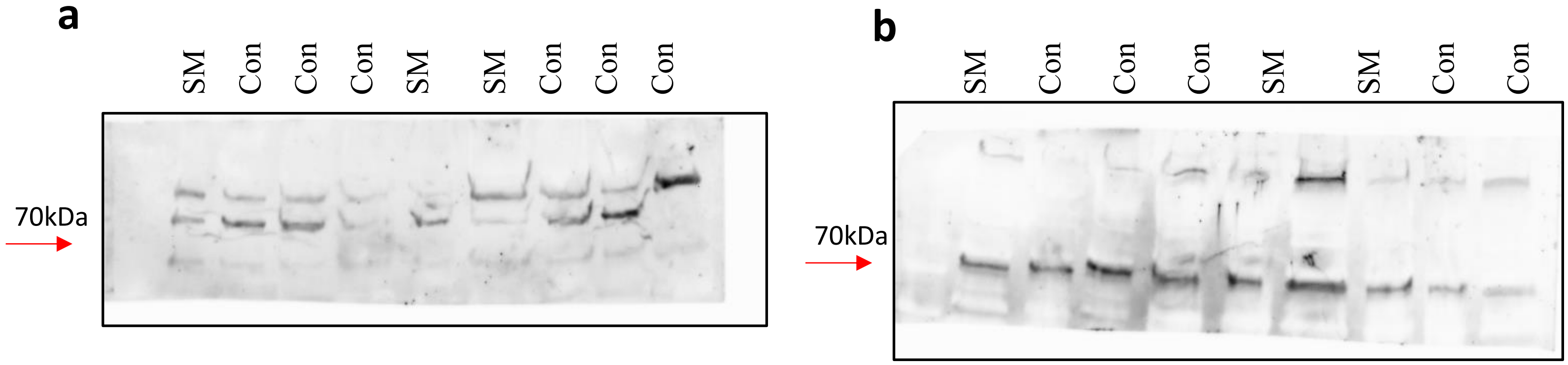


**Supplementary Figure 8.** 129S1/SvImJ 4EBP1 original western blot images. Quantification of total and phosphorylated 4EBP1 protein levels in 129S1/SvImJ mouse quadriceps. (a): total 4EBP1. SM n=3; Con n=5. (b): phosphorylated 4EBP1 (T37/46). SM n=3; Con n=6.

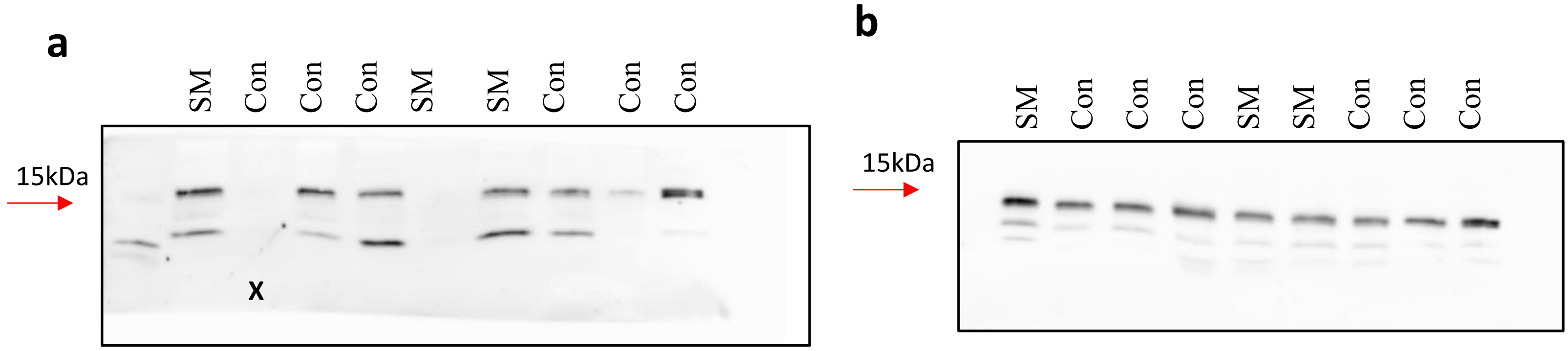

**Supplementary Figure 9.** NZO/HILtJ p70S6K1 original western blot images. Quantification of total and phosphorylated p70S6K1 protein levels in NZO/HILtJ mouse quadriceps. (a): total p70S6K1. SM n=5; Con n=6. (b): phosphorylated p70S6K1 (T-389), the final sample was run on a different gel as a repeat of lane 6. SM n=5; Con n=6.

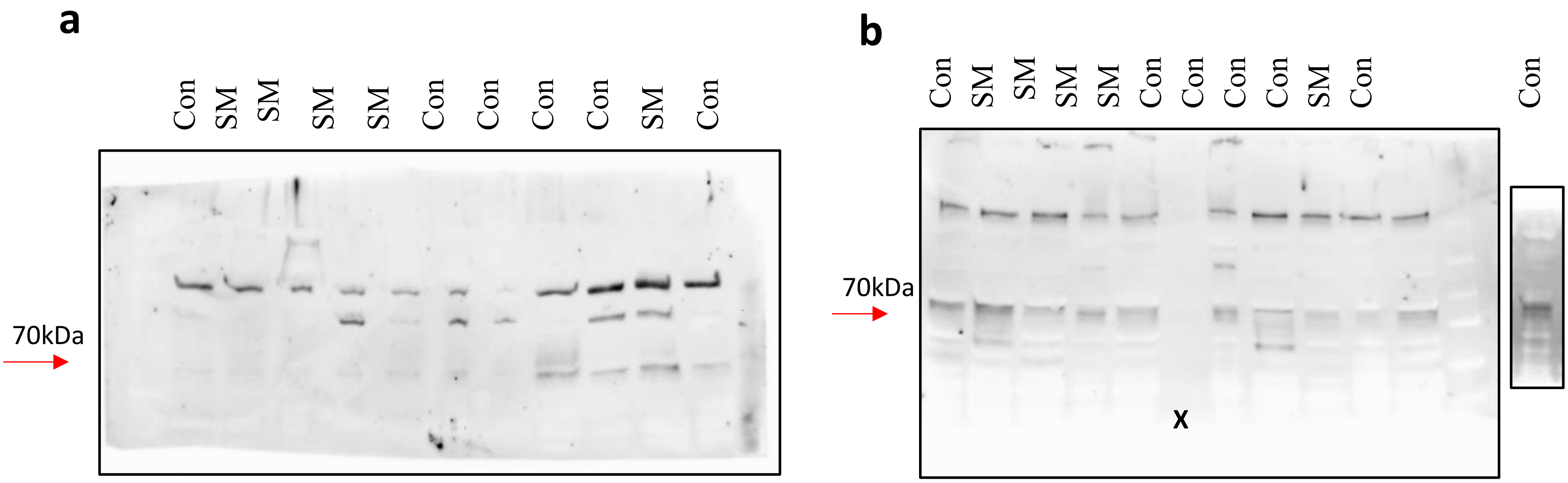


**Supplementary Figure 10.** NZO/HILtJ 4EBP1 original western blot images. Quantification of total and phosphorylated 4EBP1 protein levels in NZO/HILtJ mouse quadriceps. (a): total 4EBP1. SM n=5; Con n=5. (b): phosphorylated 4EBP1 (T37/46). SM n=5; Con n=6.

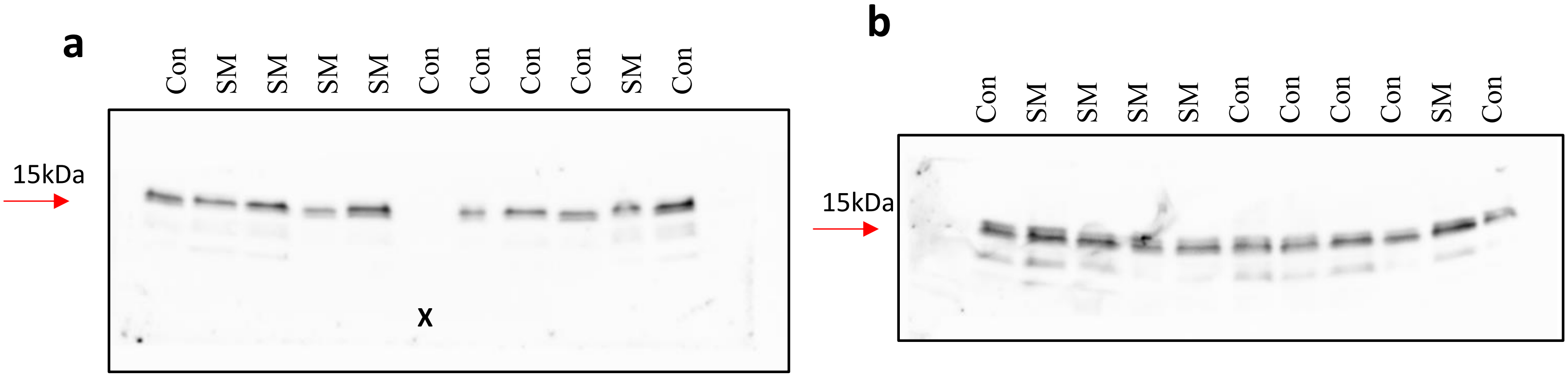

**Supplementary Figure 11.** NOD/ShiLtJ p70S6K1 original western blot images. Quantification of total and phosphorylated p70S6K1 protein levels in NOD/ShiLtJ mouse quadriceps. (a-b): total p70S6K1 ran on two gels due to space limitations. SM n=6; Con n=7. (c-d): phosphorylated p70S6K1 (T-389) ran on two gels due to space limitations. SM n=6; Con n=7.

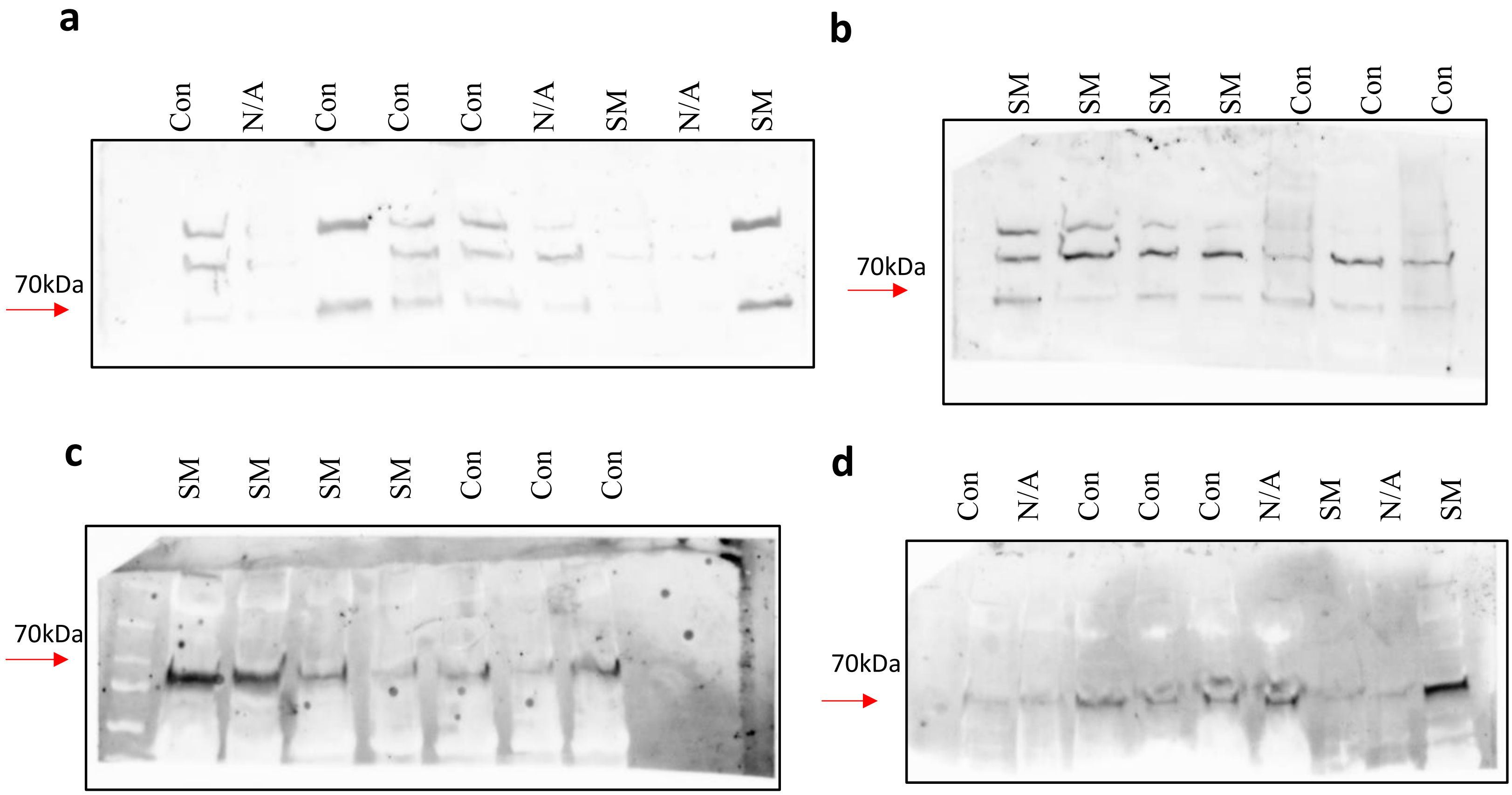


**Supplementary Figure 12.** NOD/ShiLtJ 4EBP1 original western blot images. Quantification of total and phosphorylated 4EBP1 protein levels in NOD/ShiLtJ mouse quadriceps. (a-b): total 4EBP1 ran on two gels due to space limitations. SM n=6; Con n=7. (c-d): phosphorylated 4EBP1 (T37/46) ran on two gels due to space limitations. SM n=6; Con n=7.

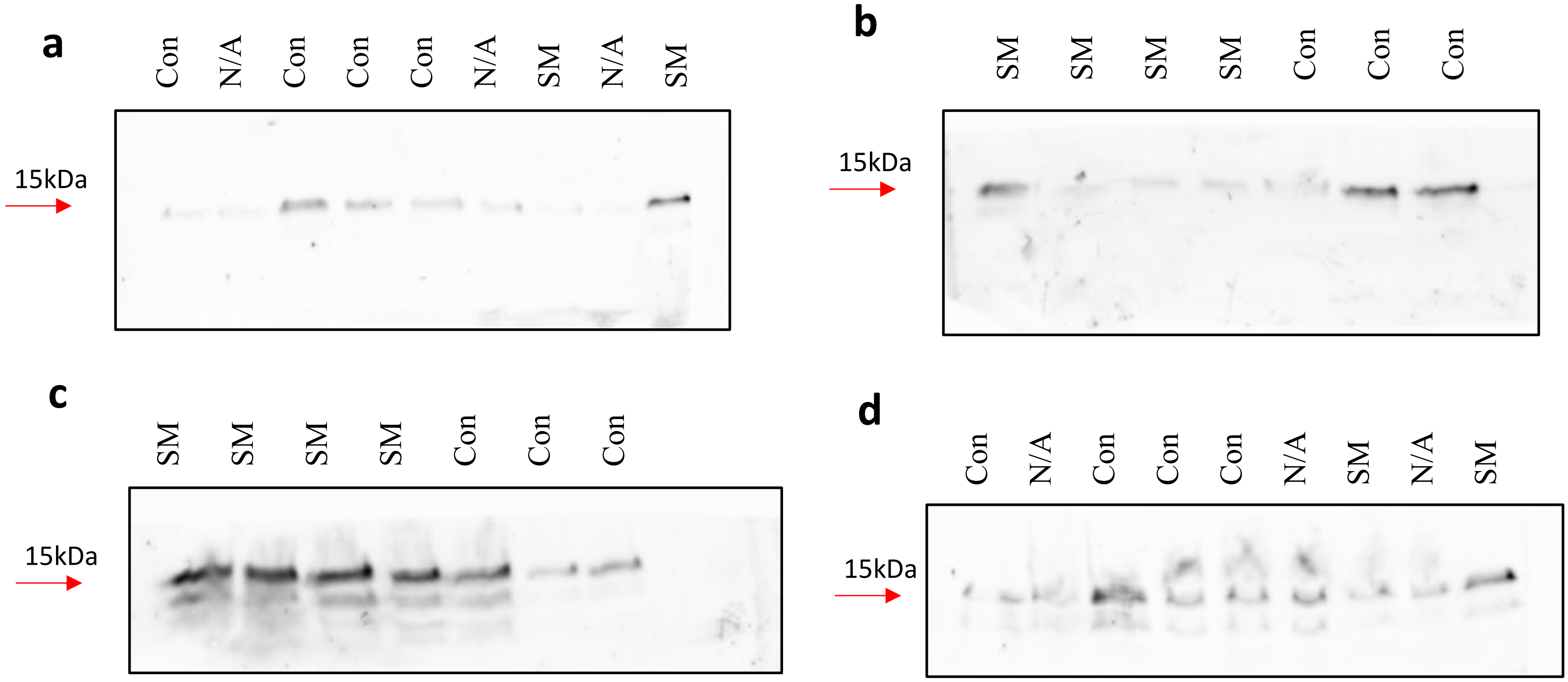

**Supplementary Figure 13.** A/J p70S6K1 original western blot images. Quantification of total and phosphorylated p70S6K1 protein levels in A/J mouse quadriceps. (a-b): total p70S6K1. The latter 4 samples were repeated due to quality issues in the first run. SM n=6; Con n=4. (b): phosphorylated p70S6K1 (T-389). SM n=8; Con n=3.

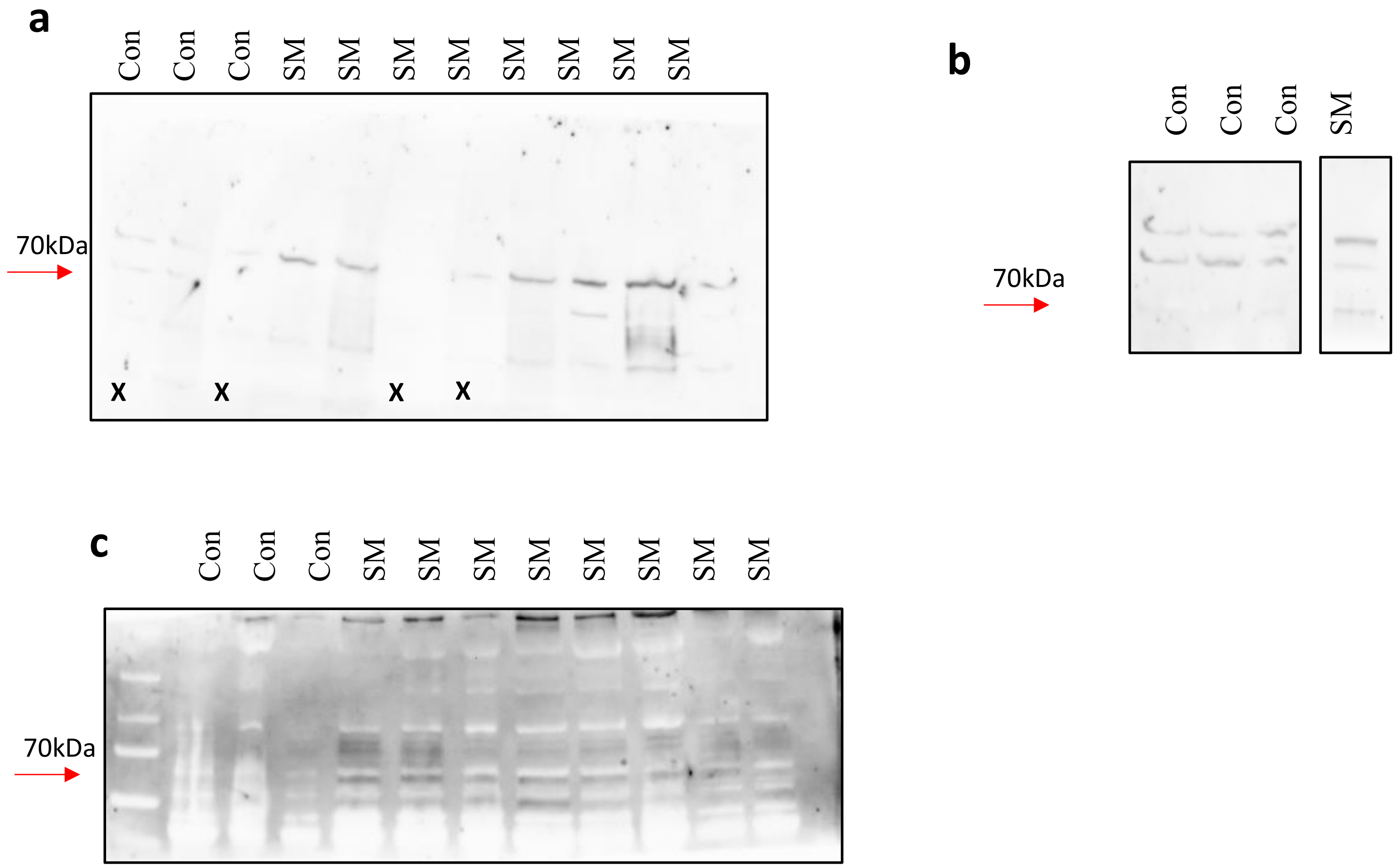


**Supplementary Figure 14.** A/J 4EBP1 original western blot images. Quantification of total and phosphorylated 4EBP1 protein levels in A/J mouse quadriceps. (a): total 4EBP1. Final lane run on separate gel due to space constraints. SM n=8; Con n=4. (b-c): phosphorylated 4EBP1 (T37/46). The latter two samples were rerun due to quality issues. SM n=7; Con n=5.

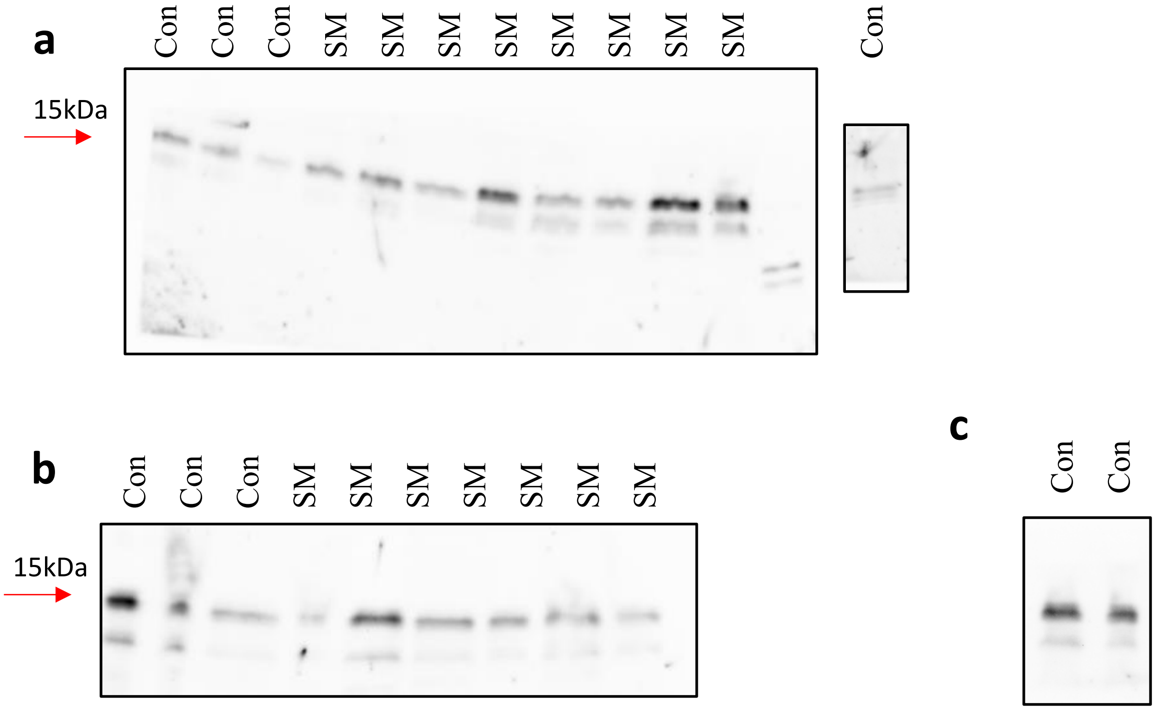

**Supplementary Figure 15.** C57BL/6J p70S6K1 original western blot images. Quantification of total and phosphorylated p70S6K1 protein levels in C57BL/6J mouse quadriceps. (a-c): total p70S6K1. Run on 3 gels due to quality issues with secondary antibodies and space limitations. SM n=8; Con n=4. (d): phosphorylated p70S6K1 (T-389). SM n=7; Con n=3.

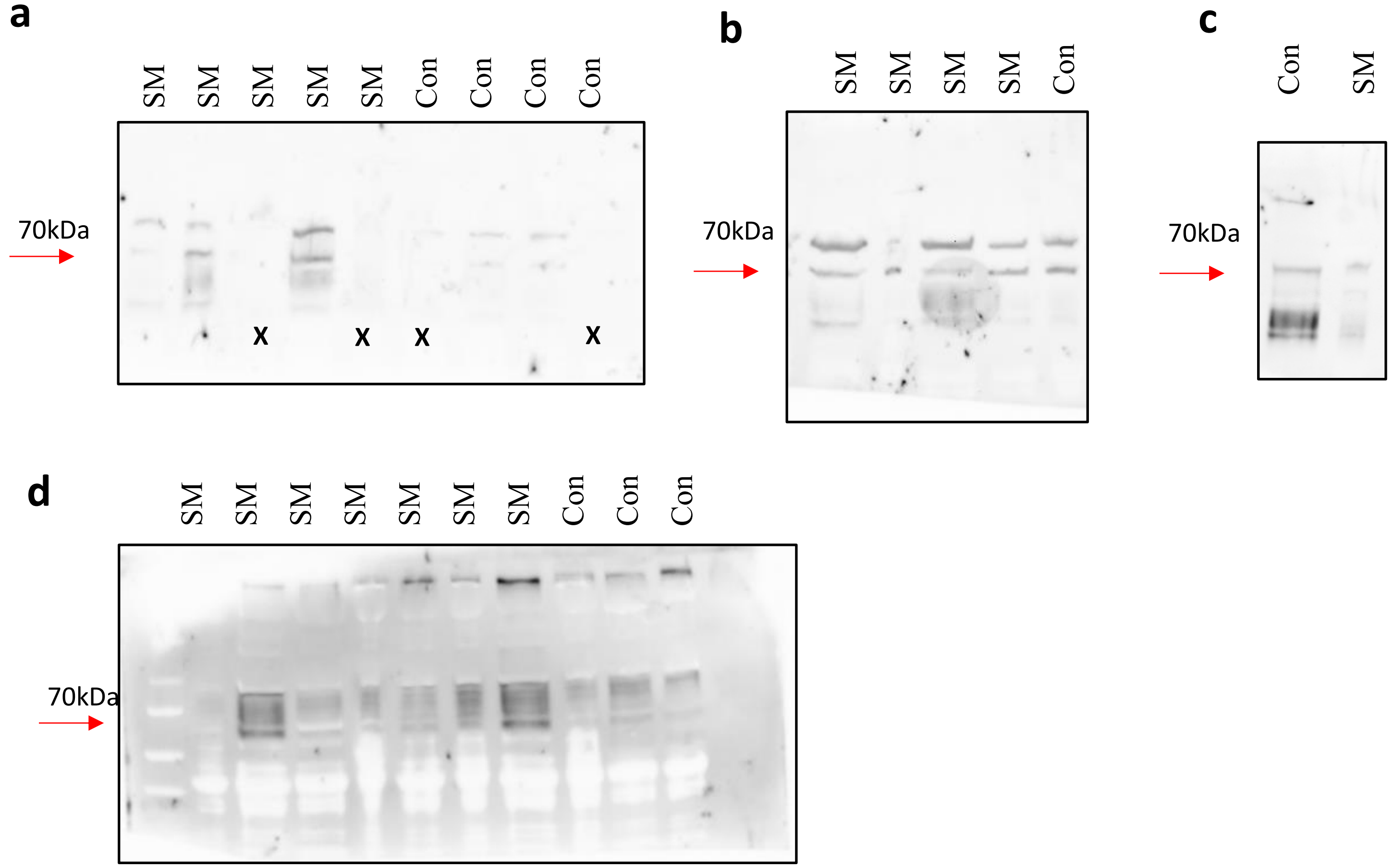


**Supplementary Figure 16.** C57BL/6J 4EBP1 original western blot images. Quantification of total and phosphorylated 4EBP1 protein levels in C57BL/6J mouse quadriceps. (a-b): total 4EBP1. Run on two separate gels due to space constraints. SM n=8; Con n=4. (c-d): phosphorylated 4EBP1 (T37/46). Run on two separate gels due to space constraints. SM n=7; Con n=5.

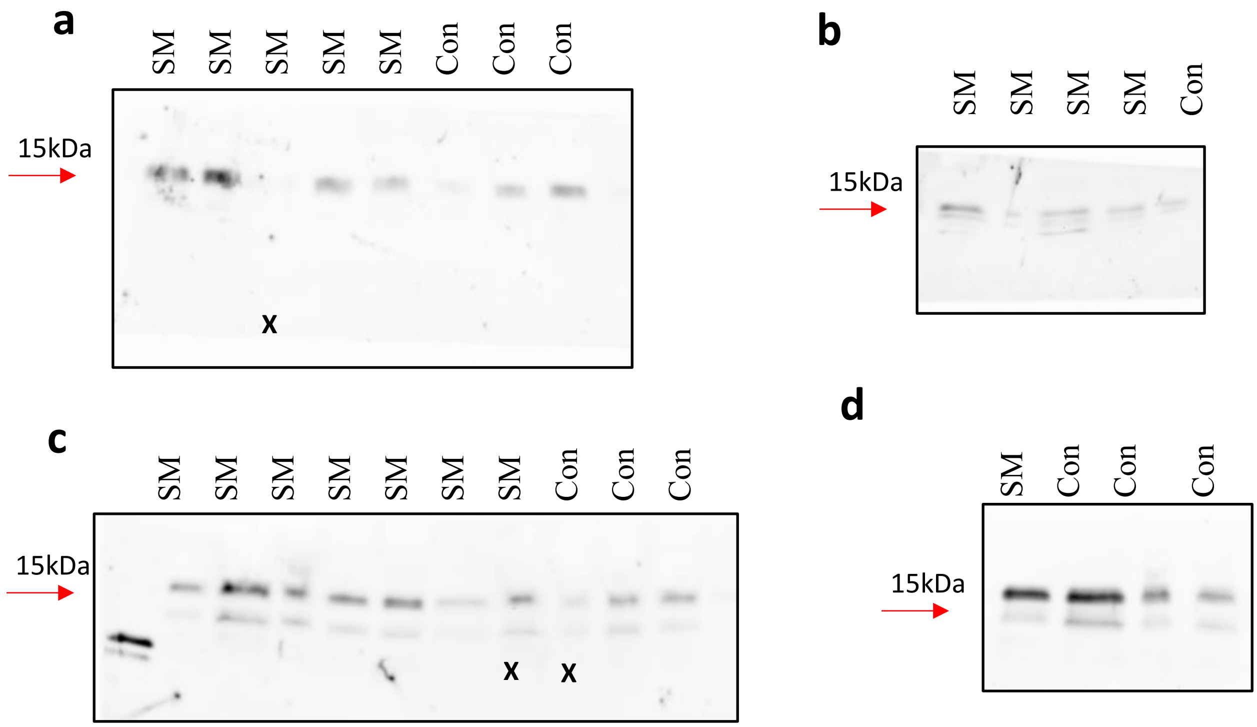
